# Hybrid assembly of an agricultural slurry virome reveals a diverse and stable community with the potential to alter the metabolism and virulence of veterinary pathogens

**DOI:** 10.1101/2020.10.08.329714

**Authors:** Ryan Cook, Steve Hooton, Urmi Trivedi, Liz King, Christine E.R. Dodd, Jon L. Hobman, Dov J. Stekel, Michael A. Jones, Andrew D. Millard

## Abstract

**Background:** Viruses are the most abundant biological entities on Earth, known to be crucial components of microbial ecosystems. However, there is little information on the viral community within agricultural waste. There are currently ^~^2.7 million dairy cattle in the UK producing 7-8% of their own bodyweight in manure daily, and 28 million tonnes annually. To avoid pollution of UK freshwaters, manure must be stored and spread in accordance with guidelines set by DEFRA. Manures are used as fertiliser, and widely spread over crop fields, yet little is known about their microbial composition. We analysed the virome of agricultural slurry over a five-month period using short and long-read sequencing.

**Results:** Hybrid sequencing uncovered more high-quality viral genomes than long or short-reads alone; yielding 7,682 vOTUs, 174 of which were complete viral genomes. The slurry virome was highly diverse and dominated by lytic bacteriophage, the majority of which represent novel genera (^~^98%). Despite constant influx and efflux of slurry, the composition and diversity of the slurry virome was extremely stable over time, with 55% of vOTUs detected in all samples over a five-month period. Functional annotation revealed a diverse and abundant range of auxiliary metabolic genes and novel features present in the community. Including the agriculturally relevant virulence factor VapE, which was widely distributed across different phage genera that were predicted to infect several hosts. Furthermore, we identified an abundance of phage-encoded diversity-generating retroelements, which were previously thought to be rare on lytic viral genomes. Additionally, we identified a group of crAssphages, including lineages that were previously thought only to be found in the human gut.

**Conclusions:** The cattle slurry virome is complex, diverse and dominated by novel genera, many of which are not recovered using long or short-reads alone. Phages were found to encode a wide range of AMGs that are not constrained to particular groups or predicted hosts, including virulence determinants and putative ARGs. The application of agricultural slurry to land may therefore be a driver of bacterial virulence and antimicrobial resistance in the environment.

## Background

Bacteriophages, or simply phages are recognised as the most abundant biological entities on the planet [1] and drive bacterial evolution through predator-prey dynamics [2,3], and horizontal gene transfer [4]. In all systems where phages have been studied in detail, they have significant ecological roles [5–7].

The contribution of phages to microbial communities has arguably been most extensively studied in the oceans [8–12]. Where, in addition to releasing large quantities of organic carbon and other nutrients through lysing bacteria, marine phages are thought to contribute to biogeochemical cycles by augmenting host metabolism with auxiliary metabolic genes (AMGs) [12–15]. Since their initial discovery, AMGs have been identified in diverse environments, including the ocean and soils [10,16]. The putative functions of AMGs are wide-ranging with the potential to alter photosynthesis, carbon metabolism, sulphur metabolism, nitrogen uptake, and complex carbohydrate metabolism [11–13,16–21].

In addition to augmenting host metabolism, phages can contribute to bacterial virulence through phage conversion via the carriage of virulence factors and toxins [22–27]. Phages have also been implicated in the transfer of antimicrobial resistance genes (ARGs) [28], however the study into the importance of phages in the transfer of ARGs has reached polarising conclusions [29,30]. Despite the vital and complex contributions of phages to microbial ecology, there is a lack of knowledge about their roles in agricultural slurry.

Manure is an unavoidable by-product from the farming of livestock. There are ^~^2.7 million dairy cattle in the UK, with ^~^1.8 million in milking herds [31]. A fully grown milking cow produces 7-8% of their own bodyweight as manure per day [32], leading to an estimated 28.31 million tonnes of manure produced by UK dairy cattle in 2010 alone [33]. These wastes are rich in nitrates and phosphates, making them valuable as a source of organic fertiliser, with an average value of £78 per cow per year [34]. However, agricultural wastes can be an environmental pollutant. Inadequate storage and agricultural run-off may lead to an increased biological oxygen demand (BOD) of freshwaters, leading to algal blooms and eutrophication [35–38]. Areas particularly at risk of nitrate pollution of ground or surface waters are classified as nitrate vulnerable zones (NVZs), and these constitute 55% of land in England [39]. For this reason, the application of organic fertilisers to fields in the UK is strictly controlled and can only be applied during certain times of the year [40]. Thus, there is the requirement to store vast volumes of slurry for several months.

To produce slurry, solids are separated from manure using apparatus such as a screw press. The liquid fraction forms the basis of slurry, which is stored in a tank or lagoon, where it is mixed with water and other agricultural wastes before its application as fertiliser. Despite being widely used as a fertiliser, the composition of the virome within slurry is poorly studied. Culture based approaches have been used to study phages infecting specific bacteria such as *E. coli* [41–43], but total viral diversity within cattle slurry remains largely unexplored.

Short-read viromics has transformed our understanding of phages in other systems, allowing an overview of the abundance and diversity of phages [8,9,12,44] and AMGs found within their genomes [12,13,16]. The power of viromics is exemplified by the study of crAssphage, which was first discovered in viromes in 2014 [45] and has subsequently been found to be the most abundant phage in the human gut and has recently been brought into culture [45–47]. However, the use of short-reads is not without limitations. Phages that contain genomic islands and/or have high micro-diversity, such as phages of the ubiquitous *Pelagibacterales* [48,49], can cause genome fragmentation during assembly [50–53]. The development of long-read sequencing technologies – most notably Pacific Biosciences (PacBio) and Oxford Nanopore Technologies (ONT) – offer a solution to such issues. The longer reads are potentially able to span the length of entire phage genomes, overcoming assembly issues resulting from repeat regions and low coverage [50–52]. The cost of longer reads is a higher error rate, which can lead to inaccurate protein prediction [54,55].

Recently, a Long-Read Linker-Amplified Shotgun Library (LASL) was developed that combines LASL library preparation with ONT MinION sequencing [56]. This approach overcame both the requirement for high DNA input for MinION sequencing and associated assembly issues with short-read sequencing. The resulting approach increased both the number and completeness of phage genomes compared to short-read assemblies [56]. An alternative approach that has utilised long-read sequencing used the ONT GridION platform to obtain entire phage genomes using an amplification-free approach on high molecular weight DNA [57]. While this approach recovered over 1,000 high quality viral genomes that could not be recovered from short-reads alone, it requires large amounts of input DNA [57], that may be inhibitory from many environments.

The aim of this work was to utilise viral metagenomics to investigate the diversity, community structure and ecological roles of viruses within dairy cattle slurry that is spread on agricultural land as an organic fertiliser.

## Methods

### DNA extraction and sequencing

DNA from the viral fraction was extracted from 10 ml of slurry as previously described [58]. Illumina sequencing was carried out on NovaSeq using 2 x 150 library. DNA from all five viral samples was pooled and subject to amplification with Illustra Ready-To-Go Genomphi V3 DNA amplification kit (GE, Healthcare) following manufacturer’s instructions. Post amplification DNA was de-branched with S1 nuclease (Thermo Fisher Scientific), following the manufacturer’s instructions and cleaned using a DNA Clean and Concentrator column (Zymo Research). Sequencing was carried by Edinburgh Genomics, with size selection of DNA to remove DNA < 5 kb prior to running on single PromethION flow cell. Reads were based called with guppy v2.3.35.

### Assembly and quality control

Illumina virome reads were trimmed with Trimmomatic v0.36 [59] using the following settings; PE illuminaclip, 2:30:10 leading:15 trailing:15 slidingwindow:4:20 minlen:50. Reads from the five samples were co-assembled with MEGAHIT v1.1.2 [60] using the settings; --k-min 21 --k-max 149 --k-step 24. Long-reads were assembled with flye v 2.6-g0d65569, reads were mapped back against the assembly with Minimap2 v2.14-r892-dirty [69] to produce BAM files and initially corrected with marginPolish v1.0.0 with ‘ allParams.np.ecoli.json’. Bacterial contamination and virus-like particle (VLP) enrichment was assessed with ViromeQC v1.0 [61].

### Identifying viral operational taxonomic units

To identify viral contigs, a number of filtering steps were applied. All contigs ≥ 10kb and circular contigs < 10kb [53] were processed using MASH v2.0 [62] separately against the RefSeq70 database [63] and a publicly available database of phage genomes (March 2020; P=0.01). If the closest RefSeq70 hit was to a phage/virus, the contig was included as a viral operational taxonomic unit (vOTU). Failing this, if the contig obtained a closer hit to the phage database than RefSeq70, the contig was included as a vOTU. Remaining contigs were included as vOTUs if they satisfied at least two of the following criteria; 1: VIBRANT v1.0.1 indicated sequence is viral [64], 2: obtained adjusted P-value ≤ 0.05 from DeepVirFinder v1.0 [65], 3: 30% of ORFs on the contig obtained a hit to a prokaryotic virus orthologous group (pVOG) [66] using Hmmscan v3.1b2 (-E 0.001) [67]. However, circular contigs < 10kb only had to satisfy either criteria 1 or 3, as DeepVirFinder scores for these contigs were inconsistent.

### Prophage analysis

A set of prophage sequences was identified from bacterial metagenomes. These were filtered as above, however contigs < 10 kb were not included even if circular. To determine which prophage vOTUs could be detected in the free viral fraction, Illumina virome reads were mapped to vOTUs using Bbmap v38.69 [68] at 90% minimum ID and the ambiguous=all flag, and Promethion reads were mapped to prophage vOTUs using Minimap2 v2.14-r892-dirty [69] with parameters “-a -x map-ont”. vOTUs were deemed as present in the free viral fraction if they obtained ≥ 1x coverage across ≥ 75% of contig length in at least one sample [53]. To determine the ends of prophages, differential coverage obtained by mapping the Illumina virome reads was investigated. Median coverage of the whole prophage was calculated and compared to median coverage across a 500 bp sliding window. If the 500 bp window had a depth of coverage ≥ 2x standard deviations lower than the median coverage of the whole prophage, this was considered a break in coverage and used to infer the ends of the prophage. An example is provided in supplementary figure 1.

To determine whether the prophage vOTUs were also assembled in the free viral assembly, contigs from the free viral fraction were mapped to prophage vOTUs using Minimap2 v2.14-r892-dirty [69] with parameters “-a -x asm10”. A prophage was considered “assembled” in the free viral fraction if a single free viral contig covered ≥ 75% of the prophage length.

### Hybrid assembly composition

Illumina reads were mapped to Promethion vOTUs using Minimap2 v2.14-r892-dirty [69] and the contigs were polished using Pilon v1.22 [70]. The Promethion vOTUs underwent multiple rounds of polishing until changes to the sequence were no longer made, or the same change was swapped back and forth between rounds of polishing. The Illumina vOTUs, hybrid vOTUs and prophage vOTUs (only those detected in the viral fraction) were de-replicated at 95% average nucleotide identity (ANI) over 80% genome length using ClusterGenomes v5.1 [71] to produce a final set of vOTUs, hereby referred to as the Final Virome. To determine assembly quality, CheckV v0.5.0 [72] was used. As this pipeline was released after the analysis in this work was performed, this was performed post-analysis.

### Alpha diversity and population dynamics

To estimate relative abundance, Illumina reads were mapped to vOTUs using Bbmap v38.69 [68] at 90% minimum ID and the ambiguous=all flag. vOTUs were deemed as present in a sample if they obtained ≥ 1x coverage across ≥ 75% of contig length [53]. The number of reads mapped to present vOTUs were normalised to reads mapped per million. Relative abundance values were analysed using Phyloseq v1.26.1 [73] in R v3.5.1 [74] to calculate diversity statistics.

Statistical testing of similarity of vOTU profiles between samples was carried out using DirtyGenes [75]. We used the randomization option with 5,000 simulations rather than chi-squared because of the small number of samples, but resampling from the null hypothesis Dirichlet distribution because there are no replicated libraries; the updated code has been uploaded to GitHub (https://github.com/LMShaw/DirtyGenes). The analysis was repeated using both the preferred cut-off of minimum 1% abundance in at least one sample and also with minimum abundance at 0.5% in at least one sample. This is because with a 1% cut-off only 7 vOTUs were included (plus an ‘other’ category binning all remaining lower abundance vOTUs) which we did not consider to be sufficiently representative; with 0.5%, 22 vOTUs were included (plus an ‘other’ category).

### Functional annotation

Final Virome vOTUs were annotated using Prokka v1.12 [76] with a custom database created from phage genome downloaded at the time (March, 2020) [77], and ORFs were compared to profile HMMs of pVOGs [66] using Hmmscan v3.1b2 (-E 0.001) [67]. Final Virome vOTU ORFs were clustered at 90% ID over 90% contig length using CD-HIT v4.6 [78] to reduce redundancy. The resultant proteins were submitted to eggNOG-mapper v2.0 [79] with default parameters, and the output was manually inspected to identify AMGs of interest. Translated ORFs identified as carbohydrate-active enzymes (CAZYmes) by eggNOG were submitted to the dbCAN2 meta-server for CAZYme identification using HMMER method to confirm their identity [16,80].

### Diversity-generating retroelement analysis

vOTUs found to encode a putative reverse transcriptase were processed using MetaCCST [81] to identify potential diversity-generating retroelements (DGRs). To identify hypervariable regions in the target gene of DGRs, reads from each sample were individually mapped to vOTUs using Bbmap v38.69 [68] at 95% minimum ID with the ambiguous=all flag. Resultant bam files were processed with Samtools v1.10 [82] to produce a mpileup file. Variants were called using VarScan v2.3 [83] mpileup2snp command with parameters “--min-coverage 10 --min-avg-qual-30”. The % SNP sites per gene were calculated for both DGR target gene(s) and all other genes on the vOTU, in order to identify if the DGR target gene(s) contained more SNP sites than on average across the vOTU.

### Taxonomy and predicted host

Final Virome vOTUs were clustered using vConTACT2 v0.9.13 [84] with parameters; --db ‘ProkaryoticViralRefSeq85-Merged’ --pcs-mode MCL --vcs-mode ClusterONE. A set of publicly available phage genome sequences (7,527), that had been deduplicated at 95% identity with dedupe.sh v36.20 [68] were included. The resultant network was visualised using Cytoscape v3.7.1 [85].

To determine if any previously known phage genomes were present in slurry viromes, reads were mapped to a dataset of publicly a set of publicly available phage genome sequences (March, 2020; 11,030), that had been deduplicated at 95% identity with dedupe.sh v36.20 [68]. Illumina reads were mapped using Bbmap v38.69 [68] at 90% minimum ID [53] and the ambiguous=all flag. Promethion reads were mapped using Minimap2 v2.14-r892-dirty [69] with parameters “-a -x map-ont”. Phages were deemed as present if they obtained ≥ 1x coverage across ≥ 75% of sequence length [53].

Putative hosts for viral vOTUs were predicted with WiSH v1.0 [86] using a database of 9,620 bacterial genomes. A p-value cut-off of 0.05 was used. Taxonomy for the predicted hosts was obtained using the R [74] package Taxonomizr v0.5.3 [87].

### Lifestyle prediction

To determine which Final Virome vOTUs were temperate, ORFs were compared to a custom set of 29 profile HMMs for transposase, integrase, excisionase, resolvase and recombinase proteins downloaded from Pfam (PF07508, PF00589, PF01609, PF03184, PF02914, PF01797, PF04986, PF00665, PF07825, PF00239, PF13009, PF16795, PF01526, PF03400, PF01610, PF03050, PF04693, PF07592, PF12762, PF13359, PF13586, PF13610, PF13612, PF13701, PF13737, PF13751, PF13808, PF13843, and PF13358) [88] using Hmmscan v3.1b2 [67] with the --cut_ga flag. Any vOTUs with an ORF which obtained a hit were classified as temperate.

### Positive selection analysis

Final Virome vOTUs which obtained ≥ 15x median coverage across ≥ 75% of contig length in every sample (excluding PHI75) were included in variant analysis. Briefly, reads were mapped onto the contigs using Bbmap v38.69 [68] at 95% minimum ID with the ambiguous=all flag, and a sorted indexed BAM file was produced. Snippy v4.4.5 [89] was used to call variants with parameters “--mapqual 0 --mincov 10”. For genes which contained at least one single nucleotide polymorphism (SNP) or multiple nucleotide polymorphism (MNP), natural selection (pN/pS) was calculated using a method adapted from Gregory *et al*. [9]. In this method, adjacent SNPs were linked as MNPs by Snippy

## Results

The farm in this study is a high-performance dairy farm in the East Midlands, UK with ^~^200 milking cattle. It houses a three million litre capacity slurry tank and an additional seven million litre lagoon to house overflow from the tank. The tank receives daily influent from the dairy farm including faeces, urine, washwater, footbath and waste milk through a slurry handling and general farm drainage system. Slurry solids are separated using a bed-press and solids are stored in a muck heap. The slurry tank and muck heap are open to the elements and the slurry tank also receives further influent from rainwater, muck heap run-off, and potentially from wildlife. The tank is emptied to ^~^10% of its maximum volume every ^~^6 weeks and the slurry is applied on fields as fertiliser.

### Comparison of short and long-read assemblies

Initial analysis of the five samples sequencing data, collected over a five month period, using viromeQC [61] indicated that one sample (PHI75) had high levels of bacterial contamination (Supplementary table 1). Sample PHI75 was excluded from further analysis, with DNA from the other four samples pooled and sequenced by Promethion sequencing.

Assembly was carried out with just Illumina or Promethion reads, resulting in 1,844 and 4,954 vOTUs ≥ 10 kb respectively. The Promethion assembly resulted in an increase in the median contig size from 12,648 bp to 14,658 compared to the Illumina only assembly (Figure 1a). The number of predicted genes per kb was also higher in the Promethion assembly. The increased error rate of Nanopore sequencing compared to Illumina sequencing is known to result in truncated gene calls [54,55]. To alleviate this, Promethion contigs were polished with Illumina reads, creating a hybrid assembly and resulting in a decrease in the number of genes per kb from 2.059 (median length: 85 aa) to 1.706 (median length: 103 aa; Figure 1b).

**Figure 1.**
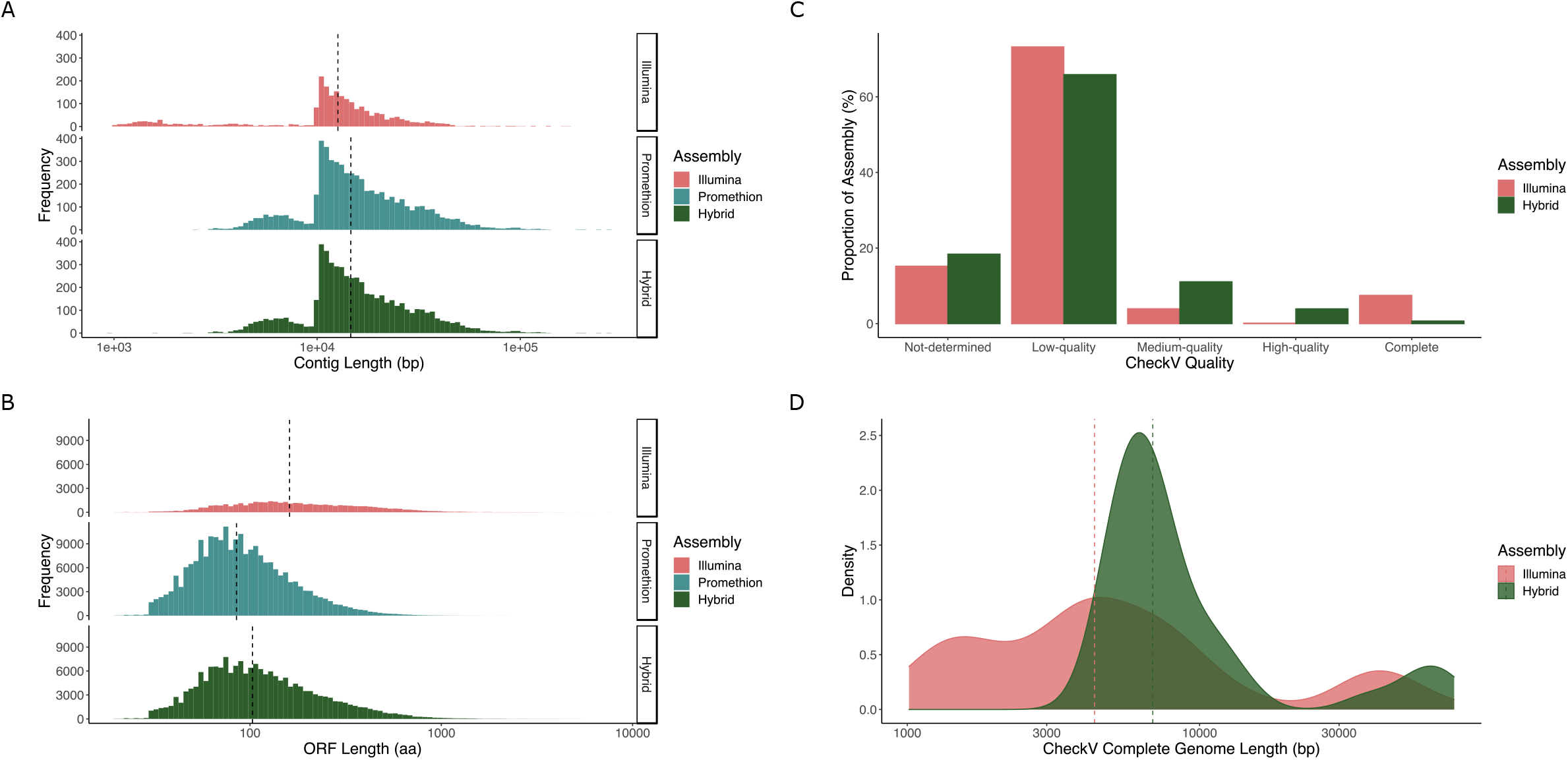
Comparison between the Illumina, Promethion and Hybrid assemblies. Histograms showing the length of vOTUs (a) and their predicted ORFs (b) within the assemblies, with the dashed line indicating median value. The different proportions of CheckV classifications of vOTUs for the Illumina and Hybrid assemblies (c) and the distribution of lengths for predicted complete genomes (d).

As whole genome amplification was used to gain sufficient material for Promethion sequencing, all diversity statistics and relative abundance data was determined from Illumina reads only. The percentage of reads that could be recruited to each different assembly was assessed. Both the Promethion (32.663%) and hybrid (33.976%) assemblies recruited more reads than the Illumina assembly (9.048%; Figure 2b). The median number of observed vOTUs per sample was higher in the Promethion (3,483) and hybrid (3,532) assemblies than that of the Illumina assembly (2,028; Figure 2a). The predicted Shannon and Simpson diversity indices increased in the hybrid (Shannon: 6.909; Simpson: 0.997) and Promethion (Shannon: 6.867; Simpson: 0.997) assemblies compared to the Illumina assembly (Shannon: 5.557; Simpson: 0.972; Figure 2c & 2d).

**Figure 2.**
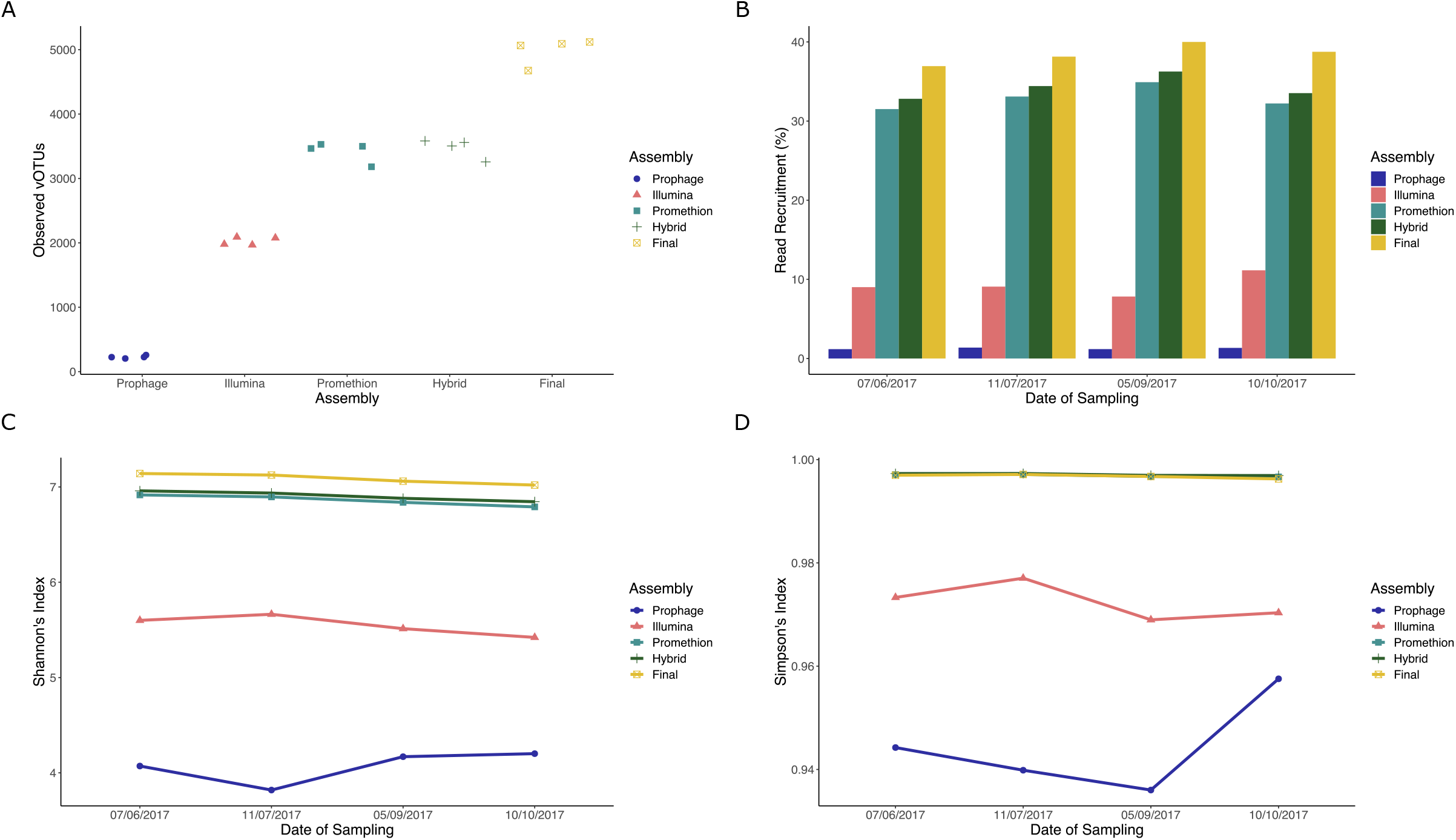
A comparison of the five different assemblies. Jitterplot shows the number of observed vOTUs per sample for the different assemblies (a). Barchart showing read recruitment over time for the different assemblies (b). Graphs showing Shanon’s (c) and Simpson’s (d) index over time for the different assemblies.

To determine the completeness and quality of the identified viral contigs, CheckV [72] was used. The hybrid assembly contained a lower proportion of low-quality genomes (65.886%), and a higher proportion of medium and high-quality (15.015%) genomes than the Illumina assembly (low-quality: 73.217%; medium and high-quality: 4.083%; Figure 1c). Conversely, the Illumina assembly contained more predicted complete genomes than the hybrid assembly (Illumina: 167; hybrid: 40). This may be due to the size selection of Promethion sequencing for longer reads, reflected in the longer average length of the complete genomes obtained from hybrid assembly (Figure 1d).

To fully understand the diversity of phages within the slurry tank, we also investigated the presence of prophage elements in the bacterial fraction. A total of 2,892 putative prophages were predicted, of which only 407 could be detected in the free phage fraction by read mapping. We combined the predicted 407 active prophages, with the Illumina and hybrid assemblies. Redundancy was removed using cluster_phages_genomes.pl [71], resulting in 7,682 vOTUs. Having established the most comprehensive virome possible, the data was further analysed.

### Characterisation of the slurry virome

The percentage of reads that could be recruited from each sample varied from 36.943% (PHI73; 07/06/2017; Figure 2b) to 39.996% (PHI76; 05/09/2017; Figure 2b). Across the five-month sampling period, the Shannon’s index alpha diversity estimates only varied from 7.02 (PHI77; 10/10/2017) to 7.141 (PHI73; 07/06/2017), suggesting a stable and diverse virome across seasons (Figure 2c & 2d). Although diverse, the virome remained stable across all sampling points with 55% (4,256) of 7,682 vOTUs found in all samples, and only 477 (^~^6%) of vOTUs unique to any one sampling point. Furthermore, testing with DirtyGenes [75] found no significant difference between the vOTU abundance profiles of the samples (P=0.1142 with 1% cut-off; P=0.863 with 0.5% cut-off). To determine if the stability in macro-diversity was mirrored by changes in micro-diversity, we assessed which predicted phage genes were under positive selection. Our analysis showed 1,610/210,997 genes (0.763%) to be under positive selection in at least one sample (Supplementary table 2). From these, putative function could be assigned to 388 translated genes. The most common predicted functions were related to phage tail (30), and phage structure (24).

To give a broader overview of the type of viruses present in the sample, pVOGs were used to infer the taxonomic classification of each vOTU. Of the vOTUs that contained proteins that matched the pVOG databases [66], 91% were associated with the order *Caudovirales*, 2.17% associated with non-tailed viruses and the remainder not classified. Approximately 10% (710) of vOTUs were identified as temperate, suggesting that the community is dominated by lytic phages of the order *Caudovirales*. The abundance of temperate vOTUs was constant across samples, ranging from 5.605% (PHI76; 05/09/2017) to 8.866% (PHI77; 10/10/2017), further demonstrating the stability of the system across time.

In order to identify the species of phages present within the slurry, all vOTUs were compared against all known phages (March, 2020) using MASH [62], with an average nucleotide identity (ANI) of > 95% as currently defined as a cut-off for phage species [90]. Only vOTUs ctg5042 and ctg217 with similarity to Mycoplasma bacteriophage L2 (accession BL2CG) and Streptococcus phage Javan630 (accession MK448997) respectively were detected. Furthermore, no vOTUs were similar to any phages that have previously been isolated from this system [41–43]. Thus, the vast majority of vOTUs represent novel phage species.

To gain an understanding of the composition at higher taxonomic levels vConTACT2 [84] was run. Only 217 (2.825%) clustered with a reference genome, indicating they are related at the genus level (Figure 3a). Notably, 18 vOTUs formed a cluster with ΦCrAss001 (accession MH675552) and phage IAS (accession KJ003983), with ctg20 appearing to be a near-complete phage genome (^~^99 kb; Figure 4b). The other 7,465 vOTUs clustered only with other vOTUs (3,369; 43.856%) or were singletons (4,096; 53.319%), indicating 5,242 putative new genera. These new genera comprised 98.037% of phages across all samples, suggesting this system is dominated by novel viruses (Figure 3b). Working on the assumption that if a vOTU within a VC was identified as temperate all other vOTUs in the cluster are, the relative abundance of temperate phages was predicted. This ranged from 13.09% (PHI76; 05/09/2017) to 16.249% (PHI77; 10/10/2017), further demonstrating the dominance of lytic viruses and stability of the system over time (Figure 3c).

**Figure 3.**
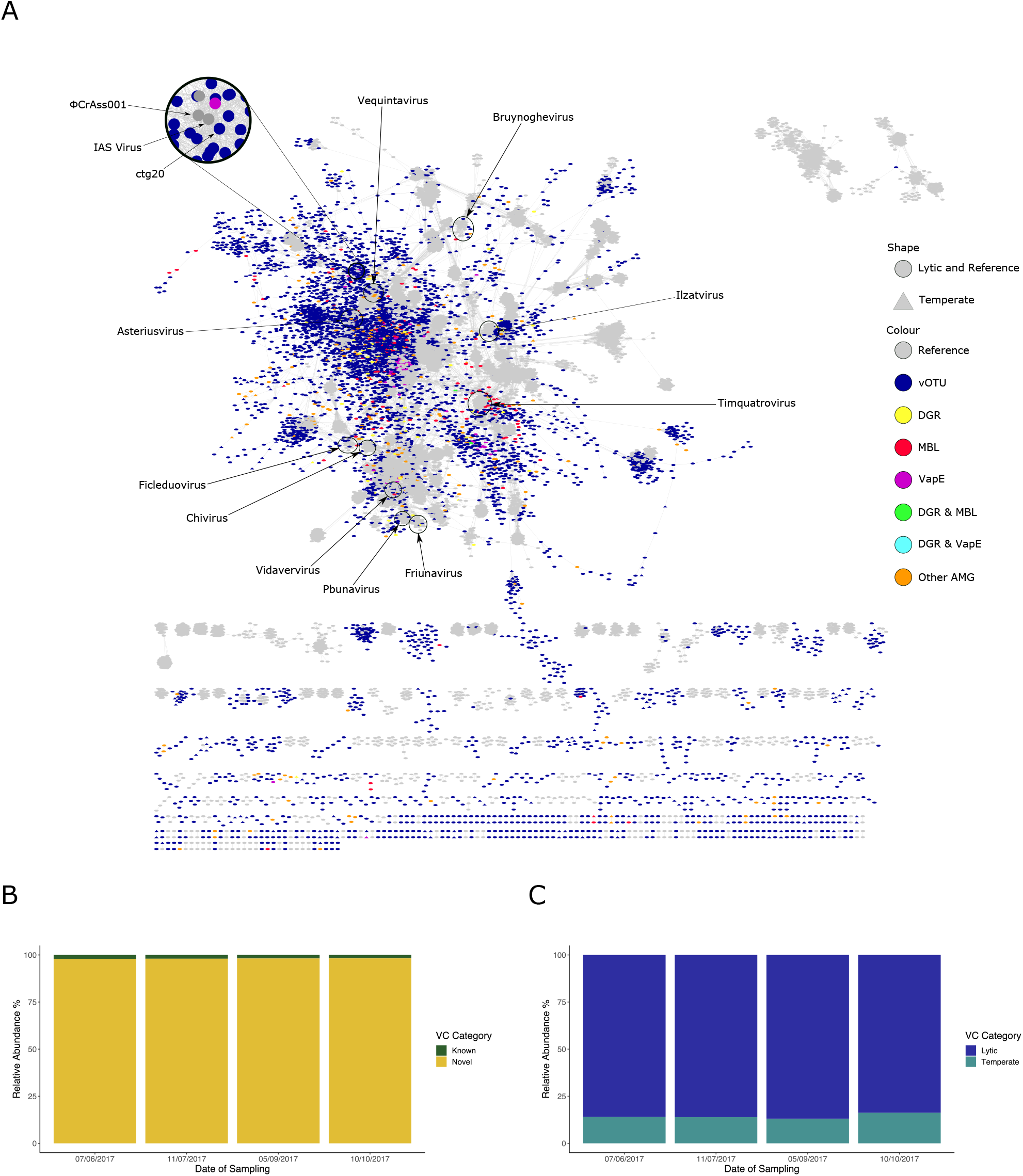
Cluster analysis of vOTUs. Network (a) shows clustering of vOTUs with reference genomes and distribution of AMGs. Barplots showing relative abundance of viral clusters based upon presence of known viruses in the same cluster (b) and their putative lifestyle (c).

**Figure 4.**
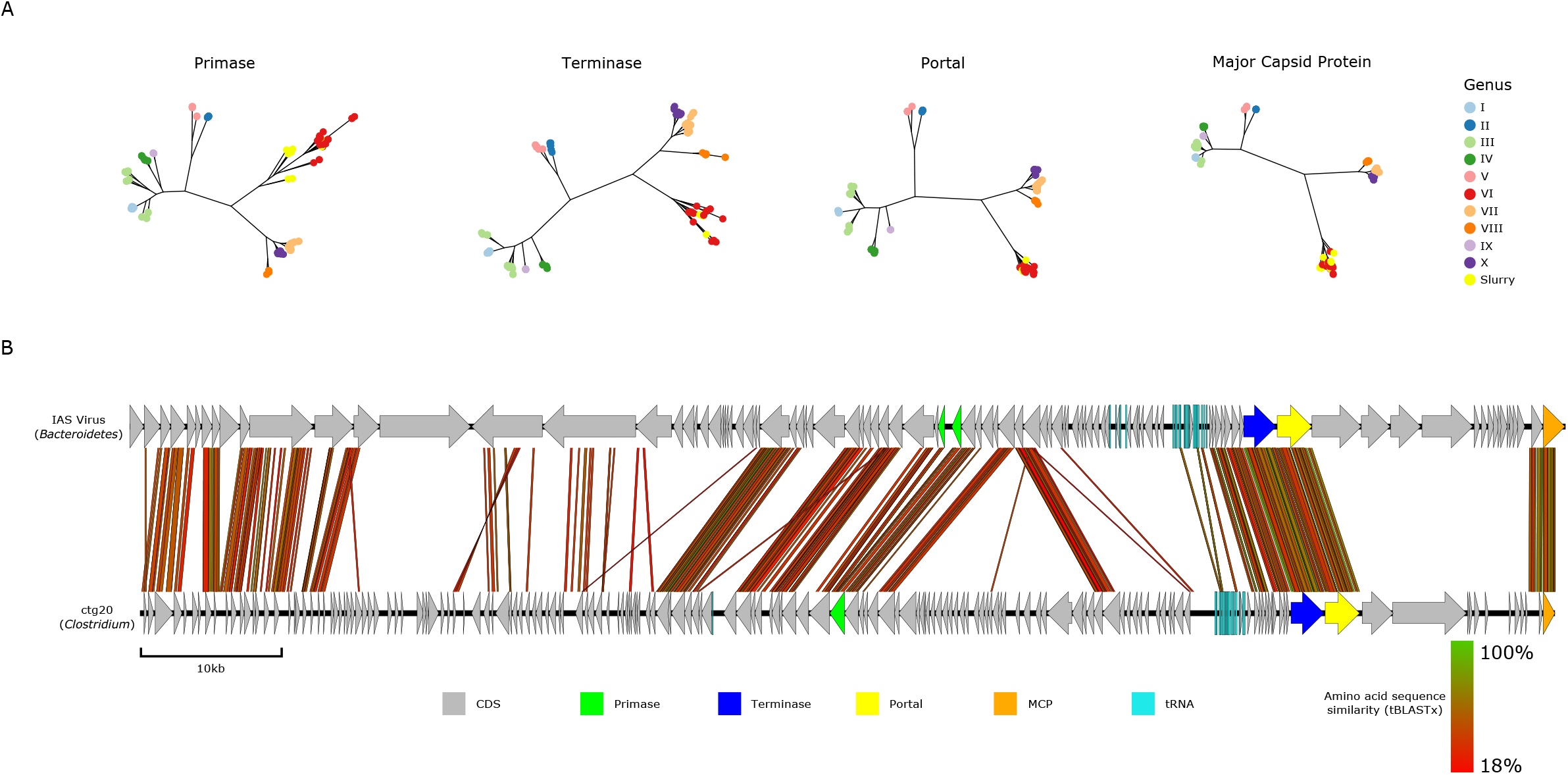
Classification of slurry crAssphages. Phylogeny of four signature genes using the method from Guerin et al. [47] shows all slurry crAssphages belong to the proposed genus VI (a). A genome comparison between the largest slurry crAssphage (ctg20) and the IAS virus, with predicted hosts shown in parentheses.

Hosts were predicted for 3,189 vOTUs and the system was found to be dominated by phages predicted to infect bacteria belonging to *Firmicutes* and *Bacteroidetes*, the most dominant phyla found in the cow gut. The proportions of host-specific abundances appeared stable across all time points (Supplementary figure 2).

### Identification of CrAss-like phages in the slurry virome

The appearance of a cluster of 18 vOTUs that are similar to crAssphage was surprising given the discovery and abundance of crAssphage in human gut viromes [45–47,91]. To further investigate this, phylogenies based on the method of Guerin *et al*. were used [47] for 15 vOTUs that contained the specific marker genes. All vOTUs formed part of the previously proposed genus VI [47], including the near complete phage (ctg20; Figure 4a; Supplementary figure 3). Furthermore, the crAssphages identified from slurry did not form a single monophyletic clade. Instead, they were interspersed with human crAssphages, with some slurry crAssphages more closely related to human crAssphages than other slurry crAssphages (Figure 4a; Supplementary figure 3). Genome comparison of ctg20 and phage IAS from genus VI identified synteny in genome architecture between the phages, yet there are clearly several areas of divergence (Figure 4b). The predicted host of ctg20 was *Clostridium*, which contrasts to the *Bacteroides and Bacteroidetes* that other crAssphages have been demonstrated or predicted to infect respectively [46,92].

### Abundance and diversity of auxiliary metabolic genes

In order to understand the role phages might have on the metabolic function of their hosts, function was assigned to proteins using eggNOG [79]. Out of 210,997 predicted proteins, only 48,819 (23.137%) could be assigned a putative function. The most abundant clusters of orthologous groups (COG) categories [93] were those associated with viral lifestyle; notably replication, recombination and repair, cell wall/membrane/envelope biogenesis, transcription and nucleotide transport and metabolism (Supplementary figure 4).

In addition to this, a number of putative AMGs were identified, including putative ARGs, CAZYmes, assimilatory sulfate reduction (ASR) genes, MazG, VapE, and Zot (Supplementary table 3). These AMGs were found to be abundant and not constrained to particular set of phages or hosts they infect (Figure 3a; Supplementary table 4). For instance, carbohydrate-active enzymes were identified on 91 vOTUs across 77 putative viral genera, with 41 vOTUs predicted to infect bacteria spanning 21 families (Supplementary table 4), and genes involved in the sulfur cycle identified on 148 vOTUs across 138 putative phage genera, with 42 vOTUs predicted to infect bacteria spanning 19 families (Supplementary table 4).

### Abundance of virulence-associated proteins

Genes encoding *zot* were identified on 36 vOTUs across 33 putative genera, predicted to infect five different families of bacteria (Supplementary table 4). The bacterial virulence factor VapE, which is found in the agricultural pathogens *Streptococcus* and *Dichelobacter*, was also detected [94–96]. VapE homologues were found on 82 vOTUs (^~^1%) across 65 clusters, including 10 high quality genomes (Figure 3a). Bacterial hosts could be predicted for 17 vOTUs and spanned 10 families of bacteria (Supplementary table 4). One vOTU (ctg217) shared ^~^95% ANI with the prophage Javan630 (accession MK448997) [97]. Genome comparison between ctg217 and Javan630 revealed highly conserved genomes, with insertion of a gene encoding a putative methyltransferase in ctg217 being the largest single difference (Figure 5).

**Figure 5.**
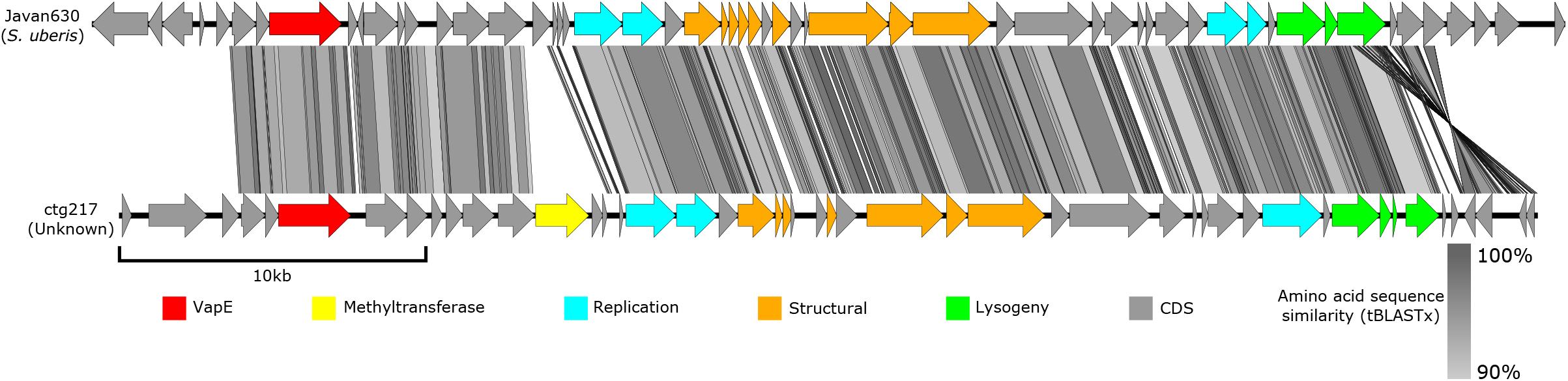
Genome comparison of the Streptococcus phage Javan630 and ctg217. A genome comparison of the VapE containing free phage (ctg217) identified in slurry and the highly similar (^~^95% ANI) Streptococcus prophage Javan630 identified from a mastitis causing strain *of Streptococcus uberis* isolated from a dairy farm in Berkshire, 2002.

### Detection of putative antimicrobial resistance genes

Putative metallo-beta-lactamases (MBLs) were identified on 146 vOTUs across 116 putative genera, with 60 vOTUs predicted to infect bacterial hosts that spanned 23 families (Supplementary table 4). Although low in sequence similarity, structural modelling with Phyre2 [98] found many of these sequences to have the same predicted structure as the novel *bla*_PNGM-1_ beta-lactamase (100% confidence over 99% coverage) [99]. Furthermore, these sequences contained conserved zinc-binding motifs characteristic of subclass B3 MBLs [99]. Phylogenetic analysis of putative phage MBLs, along with representative bacterial MBLs and a known phage-encoded *bla*_HRVM-1_ [100], showed some clustered with previously characterised bacterial MBLs and others with a characterised phage *bla*_HRVM-1_ (Supplementary figure 5). In addition to MBLs, two putative multidrug efflux pumps were identified on two vOTUs predicted to infect two different bacterial genera (Supplementary table 4).

### Identification of diversity-generating retroelements

In addition to AMGs, we also identified 202 vOTUs that carry genes encoding a reverse transcriptase. Although dsDNA phages are known to have genes that encode for a reverse transcriptase as part of diversity-generating retroelement (DGR) and the mechanism understood [101], they are rarely reported. To determine if the identified genes encoding a reverse transcriptase were part of a DGR, MetaCCST [81] was used to identify such elements. Of the 202 vOTUs carrying a reverse transcriptase gene, 82 were predicted to be part of a DGR, which accounts for ^~^1% of vOTUs in the virome. In comparison, we calculated the number of DGRs that can be identified in publicly available phage genomes (March 2020) to be 0.178%.

For vOTUS where a complete DGR system (template repeat, variable repeat, reverse transcriptase and target gene) could be identified, the most commonly predicted function of the target gene was a tail fibre. The distribution of DGRs across 74 viral clusters and 15 families of predicted host bacteria (Supplementary table 4) suggest this is not a feature that is unique to a particular VC of phages or hosts they infect (Figure 3a).

DGRs were predicted to occur on four phages that were deemed high quality complete genomes (Figure 6). These phage genomes varied in size from 40.3 kb to 52.07 kb, with two genomes containing putative integrases (k149_1459596 and k149_1764855), suggesting they are temperate, with the other two likely lytic phages (ctg154 and k149_1404499). Interestingly, phage k149_1459596 could not be detected between 07/06/2017 and 05/09/2017 but was the most abundant vOTU on 10/10/2017, representing over 3% of the viral population at that time. As vConTACT2 [84] analysis was unable to classify the phages, phylogenetic analysis was carried out with gene encoding *terL* to identify the closest known relatives (Supplementary figure 6). Phage k149_1459596 closest relative was Vibrio phage Rostov 7 (accession MK575466) and member of the *Myoviridae*. Whilst the closest known members of the three others phages are all members of the *Siphoviridae*.

**Figure 6.**
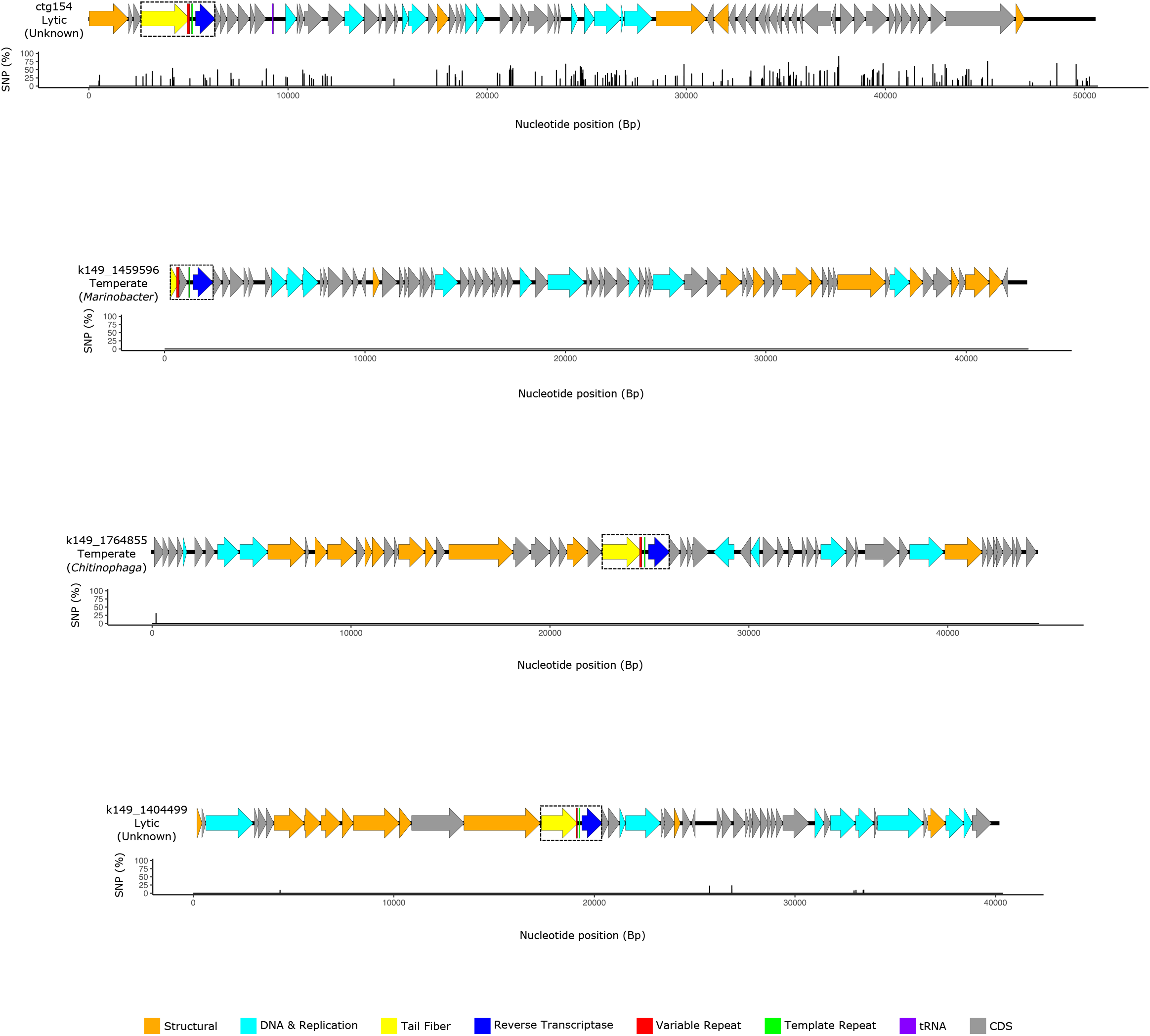
Genome maps of complete genomes with DGRs. DGR is shown in dashed box, and predicted hosts shown in parentheses. Graph beneath the genome map indicates distribution of SNP sites across the length of the genome.

We hypothesised that the widespread distribution of DGRs would reflect widespread tropism switching in these phages, and that hypervariable DGR target genes could be detected. To investigate this, we examined variants per gene and calculated which genes were under positive selection. For the 69 DGR containing vOTUs in which a target gene could be identified, 22 of these contained a higher proportion of SNP sites in the DGR target gene(s) than the average proportion of SNP sites for non-DGR target genes on that given vOTU. One of which, a predicted phage tail protein (ctg187_00023), was predicted to be under positive selection. Thus, many of the DGR target genes were more variable than other genes on a given vOTU (Figure 6).

## Discussion

### Assembly comparison

Comparison of assemblies between both short-read and long-read based sequencing methods revealed significant differences in the distribution of viral contigs and the median gene length. As has been found previously, the use of long-reads alone causes problems in gene calling due to higher error rates [54]. We therefore used short-reads to polish the long-read assembly and alleviate these issues [56]. In contrast to previous methods that used LASLs combined with ONT MinION sequencing [56], we utilised whole genome amplification followed by size selection for Promethion sequencing. Comparison of diversity statistics on Illumina, Promethion and hybrid assemblies suggest Illumina only assemblies may underestimate the diversity within a sample, whereas diversity estimates even on uncorrected Promethion assemblies is closer to that of hybrid assemblies. We also observed a number of smaller genomes that were obtained from Illumina only assemblies and were not present in the Promethion assembly. This likely results as part of the selection process for high molecular weight DNA (HMW) for Promethion sequencing that would exclude small phage genomes. Therefore, whilst long-reads improved assembly statistics, the use of long-reads alone may result in exclusion of smaller phage genomes if size selection is included. Furthermore, the inclusion of predicted prophages derived from bacterial metagenomes led to an additional 230 vOTUs that could be detected in the free viral fraction but were not assembled from virome reads. Thus, providing a more comprehensive set of viral contigs.

### Virome composition

Comparison of diversity across the period of five months revealed a highly diverse and stable virome across time. Initially this may be somewhat surprising given the dynamics of the slurry tank, which has constant inflow from animal waste, farm effluent and rainwater, and is emptied leaving only ^~^10% of the tank volume every ^~^6 weeks. We reason that most viruses in the slurry tank will originate from cow faeces, as this is the most dominant input of the tank. Host prediction suggested the virome was dominated by viruses predicted to infect bacteria belonging to *Firmicutes* and *Bacteroidetes*, which are the two most abundant bacterial phyla in the cow rumen and gut [102–104]. To date, there has been limited study into the dairy cow gut virome and its dynamics over time. However, there is a parallel with the human gut virome which is known to be temporally stable despite constant influx and efflux [105–107], and its composition influenced by environmental factors including diet [108–110]. Assuming most viruses in the slurry tank are derived from cow faeces, the controlled environment and diet of dairy cattle, results in a temporally stable virome.

Our positive selection analyses found the most common genes to be under positive selection were those involved in bacterial attachment and adsorption. We reasoned that these findings, in conjunction with the extreme stability in macro-diversity, fit with the Royal Family model of phage-host dynamics [5]. This model suggests that dominant phages are optimised to their specific ecological niche, and in the event of bacterial resistance to infection, a highly similar phage will fill that niche. Changes in community composition over time would therefore be reflected in fine-scale diversity changes, and macro-diversity would be relatively unchanged [5]. Instead of population crashes, phages may overcome bacterial resistance through positive selection of genes involved in attachment and adsorption, and are potentially accelerating the variation of these genes with DGRs.

### Diversity-generating retroelements

DGRs were first discovered in the phage BPP-1 (accession AY029185) where the reverse transcriptase, in combination with terminal repeat, produces an error prone cDNA that is then stably incorporated into the tail fibre [101]. This hypervariable region mediates the host switching of BPP-1 across different *Bordetella* species [101]. Very few DGRs have been found in cultured phage isolates since, with only two DGRs found in two temperate vibriophages [111,112]. We expanded this to 22 cultured phage isolates. Whilst not common in phage genomes, DGRs have been identified in bacterial genomes, with phage associated genes often localised next to the DGRs [112]. A recent analysis of ^~^32, 000 prophages was able to identify a further 74 DGRs in what are thought to be active prophages from diverse bacterial phyla [111]. Within this study we were able to predict a further 82 DGRs on phage genomes, four of which are thought to be complete. Two of these complete phage genomes are thought to be lytic. In fact, the majority of DGR-containing contigs in this study are thought to be lytic. Thus, demonstrating that DGRs on phage are far more common than previously found and also observed widely on lytic phages, which has not previously been observed.

Given the prevalence of DGRs, we expected to find evidence of widespread phage tropism switching by occurrence of SNPs in DGR target genes as others have done [111]. Whilst SNPs could be identified in DGR target genes supporting this, many other areas in the same phage genome contained similar levels of variation. Which is likely a result of multiple evolutionary pressures and mechanisms that are exerted on a phage genome, with DGRs only one such mechanism of creating variation.

### CrAss-like phages

Currently, crAss-like phages are classified into four subfamilies and ten genera [47], and found in a variety of environments including human waste [45–47], primate faeces [113], dog faeces [114] and termite guts [92]. Here we identified a further 18 crAss-like phages, including a near complete genome, that forms part of the proposed genus VI [47]. Genus VI is part of the *Betacrassvirinae* subfamily and currently only includes other crAss-like phages occurring within the human gut, including IAS virus that is highly abundant in HIV-1 infected individuals [115]. Thus, we have expanded the environments genus VI crAss-like phages are found in to include non-human hosts. The exact source of these phages is unknown due to the number of possible inputs of the slurry tank. However, the most likely reservoir is from cows, as this is the most abundant input. Unlike its human counterpart IAS virus, which can account for 90% of viral DNA in human faeces [45], crAss-like phages in the slurry tank were only found at low levels (^~^0.065%).

Phylogenetic analysis clearly demonstrated that human and slurry tank crAss-like phages share a common ancestor, with genetic exchange between them. The direction and route of this exchange is unclear. It may be linked to modern practices of using slurry on arable land used to produce product consumed by humans. Alternatively, it may be transferred from humans to cows via the use of biosolids derived from human waste that are applied to crops that serve as animal feed [116].

### Auxiliary metabolic genes

We identified a vast array of diverse and abundant AMGs in dairy farm slurry including putative ARGs, CAZYmes, ASR genes, MazG, VapE, and Zot. Whilst these have all been identified before in viromes from different environments [16,29,30,97,117–121], this is the first time they have been identified in slurry. The presence of different AMGs is likely a reflection of the unique composition of slurry, that has a very high water content combined with organic matter. CAZYmes were detected, which have previously been identified in viromes from mangrove soils and the cow rumen where they are thought to participate in the decomposition of organic carbon and boost host energy production during phage infection [16,122]. Given the high cellulose and hemicellulose content of slurry [123], they likely act in a similar manner within slurry to boost energy for phage replication. As well as involvement in the cycling of carbon, it also appears phage derived genes are involved in sulfur cycling within slurry. Sulfate-reducing bacteria (SRB) are active in animal wastes [124,125], and sulfate may therefore be limiting within the tank. The ASR pathway makes sulfur available for incorporation into newly synthesised molecules, such as L-cysteine and L-methionine [126], so the presence of phage encoded ASR genes on both lytic and temperate phages may overcome a metabolic bottleneck in amino acid synthesis. Alternatively, the newly synthesised ASR pathway products may be degraded for energy via the TCA cycle [127].

The AMG *mazG*, that is widespread within marine phages, in particular cyanophages [118,128,129], was also found to be abundant. The cyanophage MazG protein was originally hypothesised as a modulator of the host stringent response by altering intracellular levels of (p)ppGpp [130,131]. However, more recent work found this not to be the case [118]. The identification in a slurry tank suggests this gene is not limited to marine environments and is widespread in different phage types, although its precise role remains to be elucidated.

### Antibiotic resistance genes

There is ongoing debate as to the importance of phages in the transfer of ARGs [29,30]. We identified ARGs on ^~^ 2% of vOTUs accounting for ^~^0.082% of total predicted phage genes from assembled viral contigs. The predicted ARGs were dominated by putative MBLs, that contain core motifs and structural similarity with the known bacterial and phage MBLs *bla*_PNGM-1_ [99] and *bla*_HRVM-1_ [100] respectively. Thus, are likely functionally active, although this remains to be proven. Our estimate of the abundance of ARGs in slurry is lower than earlier reports from other environments that predict an upper estimate of ^~^0.45 % of genes in viromes are ARGs [132,133]. However, some of these studies have used unassembled reads to estimate abundance [132,133], whereas we only counted ARGs on contigs that had passed stringent filtering. Our prediction of ^~^0.082% is similar to more recent estimates of 0.001% to 0.1% in viromes from six different environments that also used assembled viromes [30], suggesting that phages might be an important reservoir of ARGs in slurry.

### Virulence-associated proteins

The virulence genes *zot* and *vapE* were found to abundant and carried by several vOTUs that were predicted to infect a range of bacterial hosts. The role of *zot* has been well studied in *Vibrio cholerae* and has previously been reported in a range of *Vibrio* and *Campylobacter* prophages [119,120,134,135]. Here, we found *zot* homologues in phages with predicted hosts other than *Vibrio* and *Camplyobacter*, further expanding the diversity of phages that carry these genes.

A similar observation was found for the virulence factor *vapE*, which has previously been found in several agricultural pathogens including *Streptococcus* and *Dichelobacter* [94–96]. *VapE* encoded on prophage elements is known to enhance the virulence *of Streptococcus* and is widespread on *Streptococcus* prophages [97]. Whilst the role of *vapE* in virulence has been established, previous work did not demonstrate the mobility of these prophage-like elements. Here we identified a high quality near-complete phage genome (ctg217) which was remarkably similar to the *vapE* encoding prophage Javan630. Phage Javan630 was originally identified as a prophage within a mastitis causing strain *of Streptococcus uberis* isolated from a dairy cow some 15 years earlier on farm ^~^100 miles away [97]. The identification of ctg217 in the free viral fraction indicates that a close relative of phage Javan630 is an active prophage. Along with the numerous other phages encoding *vapE* found in the free virome, it suggests phage are active in mediating the transfer of *vapE*. The horizontal transfer of *vapE* is of particular concern in the dairy environment where mastitis causing pathogens *Strep. uberis, Strep. agalactiae* and *Strep. dysgalactiaea* are found [136–138]. Any increase in virulence of these pathogens is detrimental to the dairy industry as it affects both animal welfare and economic viability [139]. *Streptococcus* infections result in mastitic milk, which cannot be sold and is often disposed of into slurry tanks. The continual detection of phages containing *vapE* in slurry suggests a likely continual input, given the regular emptying of the tank. The exact source of phages containing *vapE* cannot be ascertained but is likely cow faeces or mastic milk. It remains to be determined if the use of slurry as an organic fertiliser contributes to the spread of phage encoded virulence factors and toxins. However, their abundance and presence suggests it is worthy of further investigation.

## Conclusions

We have demonstrated that using a hybrid approach produces a more complete virome assembly than using short or long-reads alone. Whilst short-reads may underestimate the total viral diversity of a given environment, estimates from long-reads alone were far closer to the hybrid values than short-reads. The use of low input amplified genomic DNA allows the technique to be applied to previously sequenced metagenomes without need for further DNA extraction. We provide a comprehensive analysis of the slurry virome, demonstrating that the virome contains a diverse and stable viral community dominated by lytic viruses of novel genera. Functional annotation revealed a diverse and abundant range of AMGs including virulence factors, toxins and antibiotic resistance genes, suggesting that phages may play a significant role in mediating the transfer of these genes and augmenting both the virulence and antibiotic resistance of their hosts.

## List of Abbreviations

(p)ppGpp: Guanosine tetraphosphate or guanosine pentaphosphate
AMG: Auxiliary metabolic gene
ANI: Average nucleotide identity
ARG: Antimicrobial resistance gene
ASR: Assimilatory sulfate reduction
BOD: Biological oxygen demand
CAZYme: Carbohydrate-active enzyme
COG: Clusters of orthologous groups
DEFRA: Department for Environment, Food and Rural Affairs
DGR: Diversity-generating retroelement
eggNOG: Evolutionary genealogy of genes: Non-supervised Orthologous Groups
HGT: Horizontal gene transfer
HMW: High molecular weight
LASL: Linker-amplified shotgun library
MAG: Metagenome assembled genome
MazG: Nucleoside triphosphate pyrophosphohydrolase
MBL: Metallo-beta-lactamase
MNP: Multiple nucleotide polymorphism
ONT: Oxford Nanopore Technologies
ORF: Open reading frame
PAPS: 3⍰-phosphoadenosine-5⍰-phosphosulfate
pVOG: Prokaryotic virus orthologous groups
SNP: Single nucleotide polymorphism
SRB: Sulfate-reducing bacteria
TerL: Terminase large subunit
VapE: Virulence-associated protein E
VLP: Virus-like particle
vOTU: Viral operational taxonomic unit
Zot: *Zona occludens* toxin

## Availability of Data and Materials

Reads from Illumina and Promethion virome sequencing were submitted to the ENA under the study PRJEB38990.

## Authors’ Contributions

RC, MJ, AM, JH, CD, DS conceived the study. SH and EK carried out experiments and collected the data. RC, SH, UT and AM carried out the bioinformatic analysis. RC, MJ and AM drafted the manuscript. All authors approved and contributed to the final manuscript.

## Additional Files

## References

1. Cobián Güemes AG, Youle M, Cantú VA, Felts B, Nulton J, Rohwer F. Viruses as winners in the game of life. Annu Rev Virol. 2016;3:197–214.

2. Bohannan BJM, Lenski RE. Linking genetic change to community evolution: Insights from studies of bacteria and bacteriophage. Ecol. Lett. 2000;3:362–77.

3. Buckling A, Rainey PB. Antagonistic coevolution between a bacterium and a bacteriophage. Proc R Soc B Biol Sci. 2002;269:931–6.

4. Canchaya C, Fournous G, Chibani-Chennoufi S, Dillmann ML, Brüssow H. Phage as agents of lateral gene transfer. Curr Opin Microbiol. 2003;6:417–24.

5. Breitbart M, Bonnain C, Malki K, Sawaya NA. Phage puppet masters of the marine microbial realm. Nat. Microbiol. 2018;3:754–66.

6. Clokie MR, Millard AD, Letarov A V, Heaphy S. Phages in nature. Bacteriophage. 2011; 1:31–45.

7. Sutton TDS, Hill C. Gut bacteriophage: Current understanding and challenges. Front Endocrinol (Lausanne). 2019;10:784.

8. Paez-Espino D, Eloe-Fadrosh EA, Pavlopoulos GA, Thomas AD, Huntemann M, Mikhailova N, et al. Uncovering Earth’s virome. Nature. 2016;536:425–30.

9. Gregory AC, Zayed AA, Conceição-Neto N, Temperton B, Bolduc B, Alberti A, et al. Marine DNA viral macro- and microdiversity from Pole to Pole. Cell. 2019;177:1109–1123.

10. Hurwitz BL, U’Ren JM. Viral metabolic reprogramming in marine ecosystems. Curr Opin Microbiol. 2016;31:161–8.

11. Yooseph S, Sutton G, Rusch DB, Halpern AL, Williamson SJ, Remington K, et al. The Sorcerer II global ocean sampling expedition: Expanding the universe of protein families. PLoS Biol. 2007;5:0432–66.

12. Roux S, Brum JR, Dutilh BE, Sunagawa S, Duhaime MB, Loy A, et al. Ecogenomics and potential biogeochemical impacts of globally abundant ocean viruses. Nature. 2016;537:689–93.

13. Anantharaman K, Duhaime MB, Breier JA, Wendt KA, Toner BM, Dick GJ. Sulfur oxidation genes in diverse deep-sea viruses. Science (80-). 2014;344:757–60.

14. Zhang R, Wei W, Cai L. The fate and biogeochemical cycling of viral elements. Nat Rev Microbiol. 2014;12:850–1.

15. York A. Marine microbiology: Algal virus boosts nitrogen uptake in the ocean. Nat Rev Microbiol. 2017;15:573.

16. Jin M, Guo X, Zhang R, Qu W, Gao B, Zeng R. Diversities and potential biogeochemical impacts of mangrove soil viruses. Microbiome. 2019;7:1–15.

17. Dinsdale EA, Edwards RA, Hall D, Angly F, Breitbart M, Brulc JM, et al. Functional metagenomic profiling of nine biomes. Nature. 2008;452:629–32.

18. Sharon I, Battchikova N, Aro EM, Giglione C, Meinnel T, Glaser F, et al. Comparative metagenomics of microbial traits within oceanic viral communities. ISME J. 2011;5:1178–90.

19. Hurwitz BL, Hallam SJ, Sullivan MB. Metabolic reprogramming by viruses in the sunlit and dark ocean. Genome Biol. 2013;14:R123.

20. Hurwitz BL, Brum JR, Sullivan MB. Depth-stratified functional and taxonomic niche specialization in the “core” and “flexible” Pacific Ocean virome. ISME J. 2015;9:472–84.

21. Monier A, Chambouvet A, Milner DS, Attah V, Terrado R, Lovejoy C, et al. Host-derived viral transporter protein for nitrogen uptake in infected marine phytoplankton. Proc Natl Acad Sci U S A. 2017;114:E7489–98.

22. Freeman VJ. Studies on the virulence of bacteriophage-infected strains of *Corynebacterium diphtheriae*. J Bacteriol. 1951;61:675–88.

23. Eklund MW, Poysky FT, Meyers JA, Pelroy GA. Interspecies conversion *of Clostridium botulinum* type C to *Clostridium novyi* type A by bacteriophage. Science (80-). 1974;186:456–8.

24. Waldor MK, Mekalanos JJ. Lysogenic conversion by a filamentous phage encoding cholera toxin. Science (80-). 1996;272:1910–3.

25. Wagner PL, Livny J, Neely MN, Acheson DWK, Friedman DI, Waldor MK. Bacteriophage control of Shiga toxin 1 production and release by *Escherichia coli*. Mol Microbiol. 2002;44:957–70.

26. Khalil RKS, Skinner C, Patfield S, He X. Phage-mediated Shiga toxin (Stx) horizontal gene transfer and expression in non-Shiga toxigenic *Enterobacter* and *Escherichia coli* strains. Pathog Dis. 2016;74.

27. Fortier LC, Sekulovic O. Importance of prophages to evolution and virulence of bacterial pathogens. Virulence. 2013;4:354–65.

28. Balcázar JL. Implications of bacteriophages on the acquisition and spread of antibiotic resistance in the environment. Int. Microbiol. 2020.

29. Enault F, Briet A, Bouteille L, Roux S, Sullivan MB, Petit MA. Phages rarely encode antibiotic resistance genes: A cautionary tale for virome analyses. ISME J. 2017;11:237–47.

30. Debroas D, Siguret C. Viruses as key reservoirs of antibiotic resistance genes in the environment. ISME J. 2019;13:2856–67.

31. AHDB: UK and EU cow numbers. https://ahdb.org.uk/dairy/uk-and-eu-cow-numbers (2018). Accessed 12 Jun 2020.

32. Font-Palma C. Methods for the treatment of cattle manure—A Review. C. 2019;5:27.

33. Smith KA, Williams AG. Production and management of cattle manure in the UK and implications for land application practice. Soil Use Manag. 2016;32:73–82.

34. AHDB: Cost effective slurry storage strategies. https://dairy.ahdb.org.uk/resources-library/technical-information/health-welfare/cost-effective-slurry-storage-strategies/#.XvCQompKjwd. Accessed 12 Jun 2020.

35. De Vries JW, Groenestein CM, De Boer IJM. Environmental consequences of processing manure to produce mineral fertilizer and bio-energy. J Environ Manage. 2012;102:173–83.

36. Prapaspongsa T, Christensen P, Schmidt JH, Thrane M. LCA of comprehensive pig manure management incorporating integrated technology systems. J Clean Prod. 2010;18:1413–22.

37. Sandars DL, Audsley E, Cañete C, Cumby TR, Scotford IM, Williams AG. Environmental benefits of livestock manure management practices and technology by life cycle assessment. Biosyst Eng. 2003;84:267–81.

38. Thomassen MA, van Calker KJ, Smits MCJ, Iepema GL, de Boer IJM. Life cycle assessment of conventional and organic milk production in the Netherlands. Agric Syst. 2008;96:95–107.

39. UK Government: Nitrate vulnerable zones (NVZs). https://www.gov.uk/government/collections/nitrate-vulnerable-zones (2013). Accessed 12 Jun 2020.

40. UK Government: Use organic manures and manufactured fertilisers on farmland. https://www.gov.uk/government/publications/nitrates-and-phosphates-plan-organic-fertiliser-and-manufactured-fertiliser-use/use-organic-manures-and-manufactured-fertilisers-on-farmland. Accessed 12 Jun 2020.

41. Besler I, Sazinas P, Harrison C, Gannon L, Redgwell T, Michniewski S, et al. Genome sequence and characterization of Coliphage vB_Eco_SLUR29. PHAGE. 2020;1:38–44.

42. Sazinas P, Redgwell T, Rihtman B, Grigonyte A, Michniewski S, Scanlan DJ, et al. Comparative genomics of bacteriophage of the genus *Seuratvirus*. Genome Biol Evol. 2018;10:72–6.

43. Smith R, O’Hara M, Hobman JL, Millard AD. Draft genome sequences of 14 *Escherichia coli* phages isolated from cattle slurry. Genome Announc. 2015;3:e01364–15.

44. Brum JR, Cesar Ignacio-Espinoza J, Roux S, Doulcier G, Acinas SG, Alberti A, et al. Patterns and ecological drivers of ocean viral communities. Science (80-). 2015;348.

45. Dutilh BE, Cassman N, McNair K, Sanchez SE, Silva GGZ, Boling L, et al. A highly abundant bacteriophage discovered in the unknown sequences of human faecal metagenomes. Nat Commun. 2014;5:4498.

46. Shkoporov AN, Khokhlova E V., Fitzgerald CB, Stockdale SR, Draper LA, Ross RP, et al. ΦCrAss001 represents the most abundant bacteriophage family in the human gut and infects *Bacteroides intestinalis*. Nat Commun. 2018;9:1–8.

47. Guerin E, Shkoporov A, Stockdale SR, Clooney AG, Ryan FJ, Sutton TDS, et al. Biology and taxonomy of crAss-like bacteriophages, the most abundant virus in the human gut. Cell Host Microbe. 2018;24:653–664.e6.

48. Zhao Y, Temperton B, Thrash JC, Schwalbach MS, Vergin KL, Landry ZC, et al. Abundant SAR11 viruses in the ocean. Nature. 2013;494:357–60.

49. Martinez-Hernandez F, Fornas Ò, Lluesma Gomez M, Garcia-Heredia I, Maestre-Carballa L, López-Pérez M, et al. Single-cell genomics uncover *Pelagibacter* as the putative host of the extremely abundant uncultured 37-F6 viral population in the ocean. ISME J. 2019;13:232–6.

50. Olson ND, Treangen TJ, Hill CM, Cepeda-Espinoza V, Ghurye J, Koren S, et al. Metagenomic assembly through the lens of validation: recent advances in assessing and improving the quality of genomes assembled from metagenomes. Brief Bioinform. 2019;20:1140–50.

51. Temperton B, Giovannoni SJ. Metagenomics: Microbial diversity through a scratched lens. Curr. Opin. Microbiol. 2012;15:605–12.

52. Mizuno CM, Ghai R, Rodriguez-Valera F. Evidence for metaviromic islands in marine phages. Front Microbiol. 2014;5.

53. Roux S, Emerson JB, Eloe-Fadrosh EA, Sullivan MB. Benchmarking viromics: An *in silico* evaluation of metagenome-enabled estimates of viral community composition and diversity. PeerJ. 2017;5:e3817.

54. Watson M, Warr A. Errors in long-read assemblies can critically affect protein prediction. Nat Biotechnol. 2019;37:124–6.

55. Buck D, Weirather JL, de Cesare M, Wang Y, Piazza P, Sebastiano V, et al. Comprehensive comparison of Pacific Biosciences and Oxford Nanopore Technologies and their applications to transcriptome analysis. F1000Research. 2017;6:100.

56. Warwick-Dugdale J, Solonenko N, Moore K, Chittick L, Gregory AC, Allen MJ, et al. Long-read viral metagenomics captures abundant and microdiverse viral populations and their niche-defining genomic islands. PeerJ. 2019;7:e6800.

57. Beaulaurier J, Luo E, Eppley JM, Uyl P Den, Dai X, Burger A, et al. Assembly-free single-molecule sequencing recovers complete virus genomes from natural microbial communities. Genome Res. 2020;30:437–46.

58. Sazinas P, Michniewski S, Rihtman B, Redgwell T, Grigonyte A, Brett A, et al. Metagenomics of the viral community in three cattle slurry samples. Microbiol Resour Announc. 2019;8:e01442–18.

59. Bolger AM, Lohse M, Usadel B. Trimmomatic: a flexible trimmer for Illumina sequence data. Bioinformatics. 2014;30:2114–20.

60. Li D, Luo R, Liu CM, Leung CM, Ting HF, Sadakane K, et al. MEGAHIT v1.0: A fast and scalable metagenome assembler driven by advanced methodologies and community practices. Methods. 2016;102:3–11.

61. Zolfo M, Pinto F, Asnicar F, Manghi P, Tett A, Bushman FD, et al. Detecting contamination in viromes using ViromeQC. Nat Biotechnol. 2019;37:1408–12.

62. Ondov BD, Treangen TJ, Melsted P, Mallonee AB, Bergman NH, Koren S, et al. Mash: Fast genome and metagenome distance estimation using MinHash. Genome Biol. 2016;17:132.

63. O’Leary NA, Wright MW, Brister JR, Ciufo S, Haddad D, McVeigh R, et al. Reference sequence (RefSeq) database at NCBI: Current status, taxonomic expansion, and functional annotation. Nucleic Acids Res. 2016;44:D733–45.

64. Kieft K, Zhou Z, Anantharaman K. VIBRANT: automated recovery, annotation and curation of microbial viruses, and evaluation of viral community function from genomic sequences. Microbiome. 2020;8:90.

65. Ren J, Song K, Deng C, Ahlgren NA, Fuhrman JA, Li Y, et al. Identifying viruses from metagenomic data using deep learning. Quant Biol. 2020;8:64–77.

66. Grazziotin AL, Koonin E V, Kristensen DM. Prokaryotic Virus Orthologous Groups (pVOGs): A resource for comparative genomics and protein family annotation. Nucleic Acids Res. 2017;45:D491–8.

67. HMMER. http://hmmer.org/. Accessed 01 Mar 2020.

68. Bushnell, B: BBMap download. https://sourceforge.net/projects/bbmap/ (2013). Accessed 01 Mar 2020.

69. Li H. Minimap2: pairwise alignment for nucleotide sequences. Bioinformatics. 2018;34:3094–100.

70. Walker BJ, Abeel T, Shea T, Priest M, Abouelliel A, Sakthikumar S, et al. Pilon: An integrated tool for comprehensive microbial variant detection and genome assembly improvement. PLoS One. 2014;9:e112963.

71. Roux, S. GitHub - simroux/ClusterGenomes: Archive for ClusterGenomes scripts. https://github.com/simroux/ClusterGenomes. Accessed 01 Mar 2020.

72. Nayfach S, Pedro Camargo A, Eloe-Fadrosh E, Roux S. CheckV: assessing the quality of metagenome-assembled viral genomes. bioRxiv. 2020. https://doi.org/10.1101/2020.05.06.081778.

73. McMurdie PJ, Holmes S. Phyloseq: An R package for reproducible interactive analysis and graphics of microbiome census data. PLoS One. 2013;8:e61217.

74. R Core Team. R: A language and environment for statistical computing. Vienna: R Foundation for Statistical Computing; 2018.

75. Shaw LM, Blanchard A, Chen Q, An X, Davies P, Tötemeyer S, et al. DirtyGenes: testing for significant changes in gene or bacterial population compositions from a small number of samples. Sci Rep. 2019;9:1–10.

76. Seemann T. Prokka: Rapid prokaryotic genome annotation. Bioinformatics. 2014;30:2068–9.

77. Michniewski S, Redgwell T, Grigonyte A, Rihtman B, Aguilo-Ferretjans M, Christie-Oleza J, et al. Riding the wave of genomics to investigate aquatic coliphage diversity and activity. Environ Microbiol. 2019;21:2112–28.

78. Fu L, Niu B, Zhu Z, Wu S, Li W. CD-HIT: accelerated for clustering the next-generation sequencing data. Bioinformatics. 2012;28:3150–2.

79. Huerta-Cepas J, Szklarczyk D, Heller D, Hernández-Plaza A, Forslund SK, Cook H, et al. eggNOG 5.0: a hierarchical, functionally and phylogenetically annotated orthology resource based on 5090 organisms and 2502 viruses. Nucleic Acids Res. 2018;47:D309–14.

80. Zhang H, Yohe T, Huang L, Entwistle S, Wu P, Yang Z, et al. dbCAN2: a meta server for automated carbohydrate-active enzyme annotation. Nucleic Acids Res. 2018;46:W95–101.

81. Yan F, Yu X, Duan Z, Lu J, Jia B, Qiao Y, et al. Discovery and characterization of the evolution, variation and functions of diversity-generating retroelements using thousands of genomes and metagenomes. BMC Genomics. 2019;20:595.

82. Li H, Handsaker B, Wysoker A, Fennell T, Ruan J, Homer N, et al. The Sequence Alignment/Map format and SAMtools. Bioinformatics. 2009;25:2078–9.

83. Koboldt DC, Zhang Q, Larson DE, Shen D, McLellan MD, Lin L, et al. VarScan 2: somatic mutation and copy number alteration discovery in cancer by exome sequencing. Genome Res. 2012;22:568–76.

84. Bin Jang H, Bolduc B, Zablocki O, Kuhn JH, Roux S, Adriaenssens EM, et al. Taxonomic assignment of uncultivated prokaryotic virus genomes is enabled by gene-sharing networks. Nat Biotechnol. 2019;37:632–9.

85. Shannon P, Markiel A, Ozier O, Baliga NS, Wang JT, Ramage D, et al. Cytoscape: A software environment for integrated models of biomolecular interaction networks. Genome Res. 2003;13:2498–504.

86. Galiez C, Siebert M, Enault F, Vincent J, Söding J. WIsH: who is the host? Predicting prokaryotic hosts from metagenomic phage contigs. Bioinformatics. 2017;33:3113–4.

87. Sherrill-Mix, S: taxonomizr: Functions to Work with NCBI Accessions and Taxonomy. https://cran.r-project.org/web/packages/taxonomizr/ (2018). Accessed 01 Mar 2020.

88. El-Gebali S, Mistry J, Bateman A, Eddy SR, Luciani A, Potter SC, et al. The Pfam protein families database in 2019. Nucleic Acids Res. 2019;47:D427–32.

89. Seemann, T: snippy: Rapid haploid variant calling and core genome alignment. https://github.com/tseemann/snippy (2015). Accessed 01 Mar 2020.

90. Adriaenssens EM, Rodney Brister J. How to name and classify your phage: An informal guide. Viruses. 2017;9:1–9.

91. Shkoporov AN, Clooney AG, Sutton TDS, Ryan FJ, Daly KM, Nolan JA, et al. The human gut virome is highly diverse, stable, and individual specific. Cell Host Microbe. 2019;26:527–541.

92. Yutin N, Makarova KS, Gussow AB, Krupovic M, Segall A, Edwards RA, et al. Discovery of an expansive bacteriophage family that includes the most abundant viruses from the human gut. Nat Microbiol. 2018;3:38–46.

93. Tatusov RL, Galperin MY, Natale DA, Koonin E V. The COG database: a tool for genome-scale analysis of protein functions and evolution. Nucleic Acids Res. 2000;28:33–6.

94. Billington SJ, Johnston JL, Rood JI. Virulence regions and virulence factors of the ovine footrot pathogen, *Dichelobacter nodosus*. FEMS Microbiol Lett. 1996;145:147–56.

95. Bloomfield GA, Whittle G, McDonagh MB, Katz ME, Cheetham BF. Analysis of sequences flanking the vap regions *of Dichelobacter nodosus*: Evidence for multiple integration events, a killer system, and a new genetic element. Microbiology. 1997;143:553–62.

96. Ji X, Sun Y, Liu J, Zhu L, Guo X, Lang X, et al. A novel virulence-associated protein, *vapE*, in *Streptococcus suis* serotype 2. Mol Med Rep. 2016;13:2871–7.

97. Rezaei Javan R, Ramos-Sevillano E, Akter A, Brown J, Brueggemann AB. Prophages and satellite prophages are widespread in *Streptococcus* and may play a role in pneumococcal pathogenesis. Nat Commun. 2019;10:1–14.

98. Kelley LA, Mezulis S, Yates CM, Wass MN, Sternberg MJE. The Phyre2 web portal for protein modeling, prediction and analysis. Nat Protoc. 2015;10:845–58.

99. Park KS, Kim TY, Kim JH, Lee JH, Jeon JH, Karim AM, et al. PNGM-1, a novel subclass B3 metallo-β-lactamase from a deep-sea sediment metagenome. J Glob Antimicrob Resist. 2018;14:302–5.

100. Moon K, Jeon JH, Kang I, Park KS, Lee K, Cha CJ, et al. Freshwater viral metagenome reveals novel and functional phage-borne antibiotic resistance genes. Microbiome. 2020;8:75.

101. Liu M, Deora R, Doulatov SR, Gingery M, Eiserling FA, Preston A, et al. Reverse transcriptase-mediated tropism switching in *Bordetella* bacteriophage. Science (80-). 2002;295:2091–4.

102. Kim M, Wells JE. A meta-analysis of bacterial diversity in the feces of cattle. Curr Microbiol. 2016;72:145–51.

103. Delgado B, Bach A, Guasch I, González C, Elcoso G, Pryce JE, et al. Whole rumen metagenome sequencing allows classifying and predicting feed efficiency and intake levels in cattle. Sci Rep. 2019;9:1–13.

104. Li F, Hitch TCA, Chen Y, Creevey CJ, Guan LL. Comparative metagenomic and metatranscriptomic analyses reveal the breed effect on the rumen microbiome and its associations with feed efficiency in beef cattle. Microbiome. 2019;7:6.

105. Garmaeva S, Sinha T, Kurilshikov A, Fu J, Wijmenga C, Zhernakova A. Studying the gut virome in the metagenomic era: Challenges and perspectives. BMC Biol. 2019;17:1–14.

106. Reyes A, Haynes M, Hanson N, Angly FE, Heath AC, Rohwer F, et al. Viruses in the faecal microbiota of monozygotic twins and their mothers. Nature. 2010;466:334–8.

107. Minot S, Bryson A, Chehoud C, Wu GD, Lewis JD, Bushman FD. Rapid evolution of the human gut virome. Proc Natl Acad Sci U S A. 2013;110:12450–5.

108. Minot S, Sinha R, Chen J, Li H, Keilbaugh SA, Wu GD, et al. The human gut virome: Inter-individual variation and dynamic response to diet. Genome Res. 2011;21:1616–25.

109. Lim ES, Zhou Y, Zhao G, Bauer IK, Droit L, Ndao IM, et al. Early life dynamics of the human gut virome and bacterial microbiome in infants. Nat Med. 2015;21:1228–34.

110. Moreno-Gallego JL, Chou SP, Di Rienzi SC, Goodrich JK, Spector TD, Bell JT, et al. Virome diversity correlates with intestinal microbiome diversity in adult monozygotic twins. Cell Host Microbe. 2019;25:261–272.

111. Benler S, Cobián-Güemes AG, McNair K, Hung SH, Levi K, Edwards R, et al. A diversity-generating retroelement encoded by a globally ubiquitous *Bacteroides* phage 06 Biological Sciences 0605 Microbiology. Microbiome. 2018;6:1–10.

112. Wu L, Gingery M, Abebe M, Arambula D, Czornyj E, Handa S, et al. Diversity-generating retroelements: Natural variation, classification and evolution inferred from a large-scale genomic survey. Nucleic Acids Res. 2018;46:11–24.

113. Edwards RA, Vega AA, Norman HM, Ohaeri M, Levi K, Dinsdale EA, et al. Global phylogeography and ancient evolution of the widespread human gut virus crAssphage. Nat Microbiol. 2019;4:1727–36.

114. Cuscó A, Salas A, Torre C, Francino O. Shallow metagenomics with Nanopore sequencing in canine fecal microbiota improved bacterial taxonomy and identified an uncultured CrAssphage. bioRxiv. 2019. https://doi.org/10.1101/585067.

115. Oude Munnink BB, Canuti M, Deijs M, de Vries M, Jebbink MF, Rebers S, et al. Unexplained diarrhoea in HIV-1 infected individuals. BMC Infect Dis. 2014;14:22.

116. Biosolids Assurance Scheme: About biolsolids: Assured biosolids. https://assuredbiosolids.co.uk/about-biosolids/ (2020). Accessed 12 Jun 2020.

117. Gao SM, Schippers A, Chen N, Yuan Y, Zhang MM, Li Q, et al. Depth-related variability in viral communities in highly stratified sulfidic mine tailings. Microbiome. 2020;8:89.

118. Rihtman B, Bowman-Grahl S, Millard A, Corrigan RM, Clokie MRJ, Scanlan DJ. Cyanophage MazG is a pyrophosphohydrolase but unable to hydrolyse magic spot nucleotides. Environ Microbiol Rep. 2019;11:448–55.

119. Liu F, Lee H, Lan R, Zhang L. Zonula occludens toxins and their prophages in *Campylobacter* species. Gut Pathog. 2016;8:43.

120. Castillo D, Pérez-Reytor D, Plaza N, Ramírez-Araya S, Blondel CJ, Corsini G, et al. Exploring the genomic traits of non-toxigenic *Vibrio parahaemolyticus* strains isolated in southern Chile. Front Microbiol. 2018;9:161.

121. Romero P, Croucher NJ, Hiller NL, Hu FZ, Ehrlich GD, Bentley SD, et al. Comparative genomic analysis of ten *Streptococcus pneumoniae* temperate bacteriophages. J Bacteriol. 2009;191:4854–62.

122. Anderson CL, Sullivan MB, Fernando SC. Dietary energy drives the dynamic response of bovine rumen viral communities. Microbiome. 2017;5:155.

123. Chen S, Liao W, C L, Wen Z, Kincaid RL, Harrison JH, et al. Value-added chemicals from animal manure. Pacific Northwest Natl Lab. 2003;PNNL-14495:1–142.

124. Cook KL, Whitehead TR, Spence C, Cotta MA. Evaluation of the sulfate-reducing bacterial population associated with stored swine slurry. Anaerobe. 2008; 14:172–80.

125. St-Pierre B, Wright ADG. Implications from distinct sulfate-reducing bacteria populations between cattle manure and digestate in the elucidation of H2S production during anaerobic digestion of animal slurry. Appl Microbiol Biotechnol. 2017;101:5543–56.

126. Rückert C. Sulfate reduction in microorganisms — recent advances and biotechnological applications. Curr. Opin. Microbiol. 2016;33:140–6.

127. Howard-Varona C, Lindback MM, Bastien GE, Solonenko N, Zayed AA, Jang H Bin, et al. Phage-specific metabolic reprogramming of virocells. ISME J. 2020;14:881–95.

128. Sullivan MB, Huang KH, Ignacio-Espinoza JC, Berlin AM, Kelly L, Weigele PR, et al. Genomic analysis of oceanic cyanobacterial myoviruses compared with T4-like myoviruses from diverse hosts and environments. Environ Microbiol. 2010;12:3035–56.

129. Millard AD, Zwirglmaier K, Downey MJ, Mann NH, Scanlan DJ. Comparative genomics of marine cyanomyoviruses reveals the widespread occurrence of *Synechococcus* host genes localized to a hyperplastic region: Implications for mechanisms of cyanophage evolution. Environ Microbiol. 2009;11:2370–87.

130. Clokie MRJ, Mann NH. Marine cyanophages and light. Environ Microbiol. 2006;8:2074–82.

131. Clokie MRJ, Millard AD, Mann NH. T4 genes in the marine ecosystem: studies of the T4-like cyanophages and their role in marine ecology. Virol J. 2010;7:291.

132. Balcazar JL. Bacteriophages as vehicles for antibiotic resistance genes in the environment. PLoS Pathog. 2014;10:e1004219–e1004219.

133. Lekunberri I, Subirats J, Borrego CM, Balcázar JL. Exploring the contribution of bacteriophages to antibiotic resistance. Environ Pollut. 2017;220:981–4.

134. Koonin E V. The second cholera toxin, Zot, and its plasmid-encoded and phage-encoded homologues constitute a group of putative ATPases with an altered purine NTP-binding motif. FEBS Lett. 1992;312:3–6.

135. Schmidt E, Kelly SM, van der Walle CF. Tight junction modulation and biochemical characterisation of the zonula occludens toxin C-and N-termini. FEBS Lett. 2007;581:2974–80.

136. Zadoks RN, Middleton JR, McDougall S, Katholm J, Schukken YH. Molecular epidemiology of mastitis pathogens of dairy cattle and comparative relevance to humans. J Mammary Gland Biol Neoplasia. 2011;16:357–72.

137. Whist AC, Østerås O, Sølverød L. *Streptococcus dysgalactiae* isolates at calving and lactation performance within the same lactation. J Dairy Sci. 2007;90:766–78.

138. Keefe GP. *Streptococcus agalactiae* mastitis: A review. Can Vet J. 1997;38:429–37.

139. AHDB: Mastitis in dairy cows. https://dairy.ahdb.org.uk/technical-information/animal-health-welfare/mastitis/#.Xt5XWZ5Kjwc (2018). Accessed 12 Jun 2020.

