## Supplementary figure for "Hybrid assembly of an agricultural slurry virome reveals a diverse and stable community with the potential to alter the metabolism and virulence of veterinary pathogens"

Supplementary figure 1

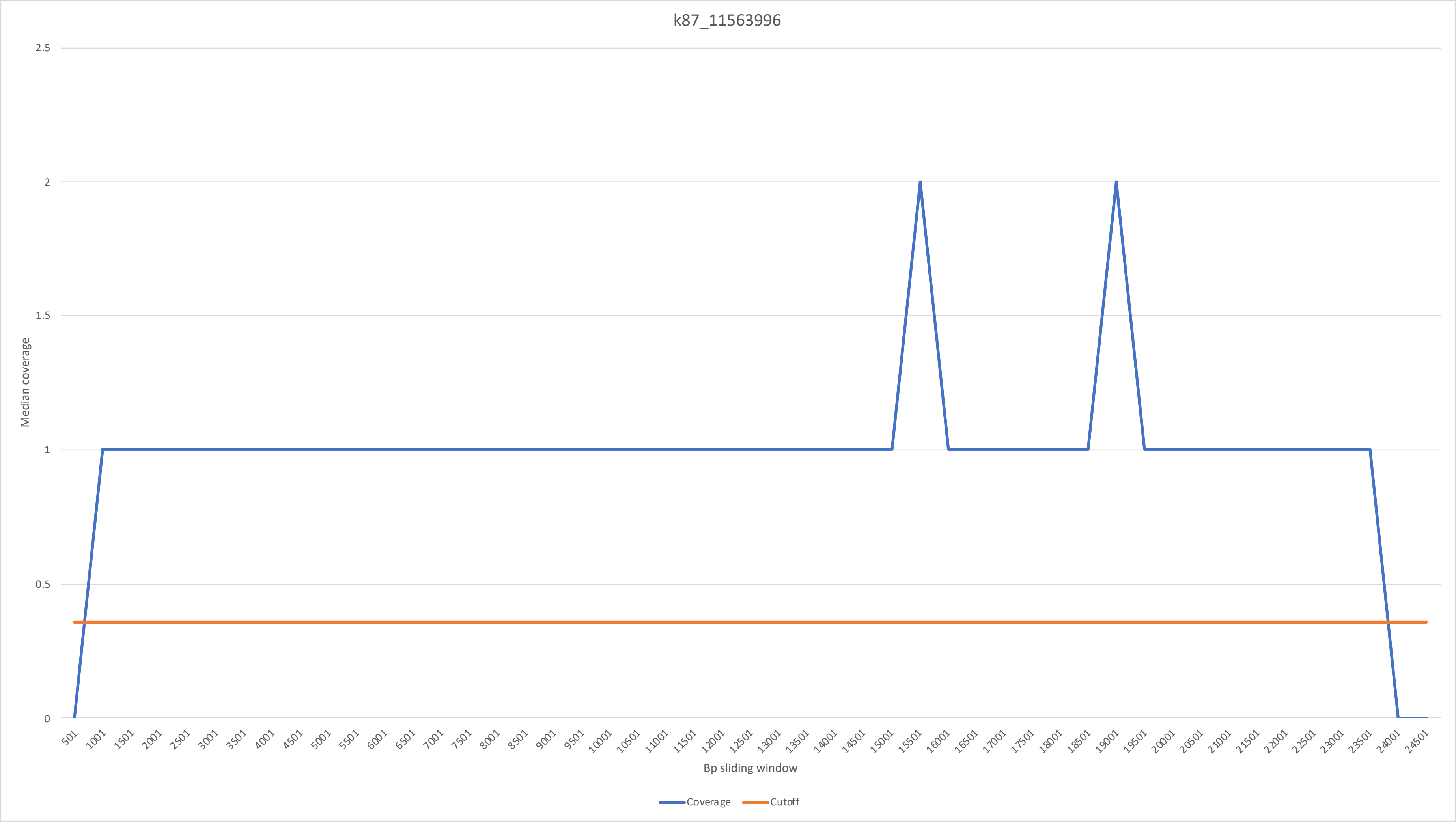

Supplementary figure 2

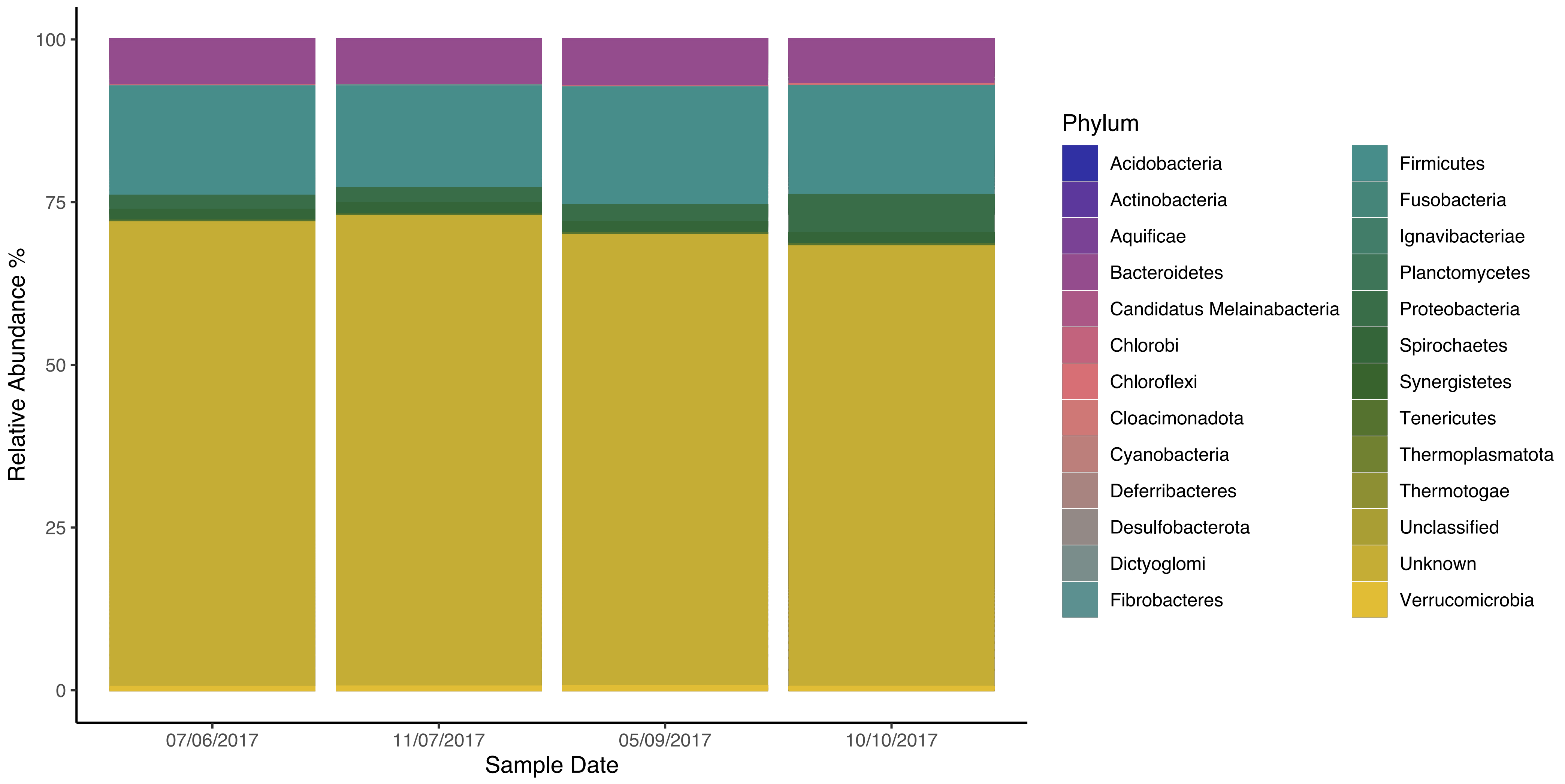

### Supplementary figure 3

Primase

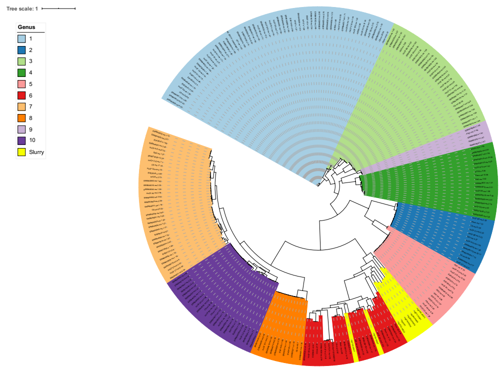


Terminase

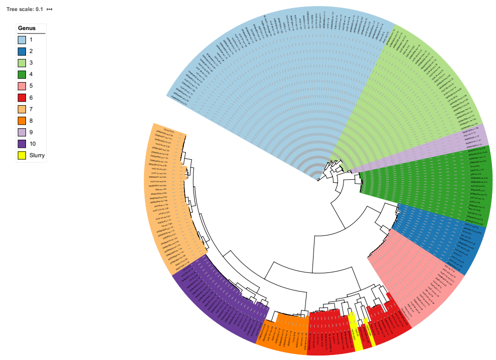


Portal

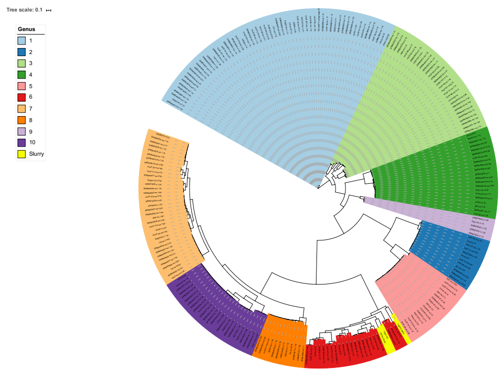


Major Capsid Protein

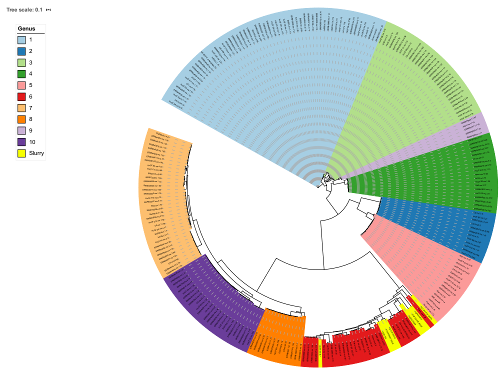

Supplementary figure 4

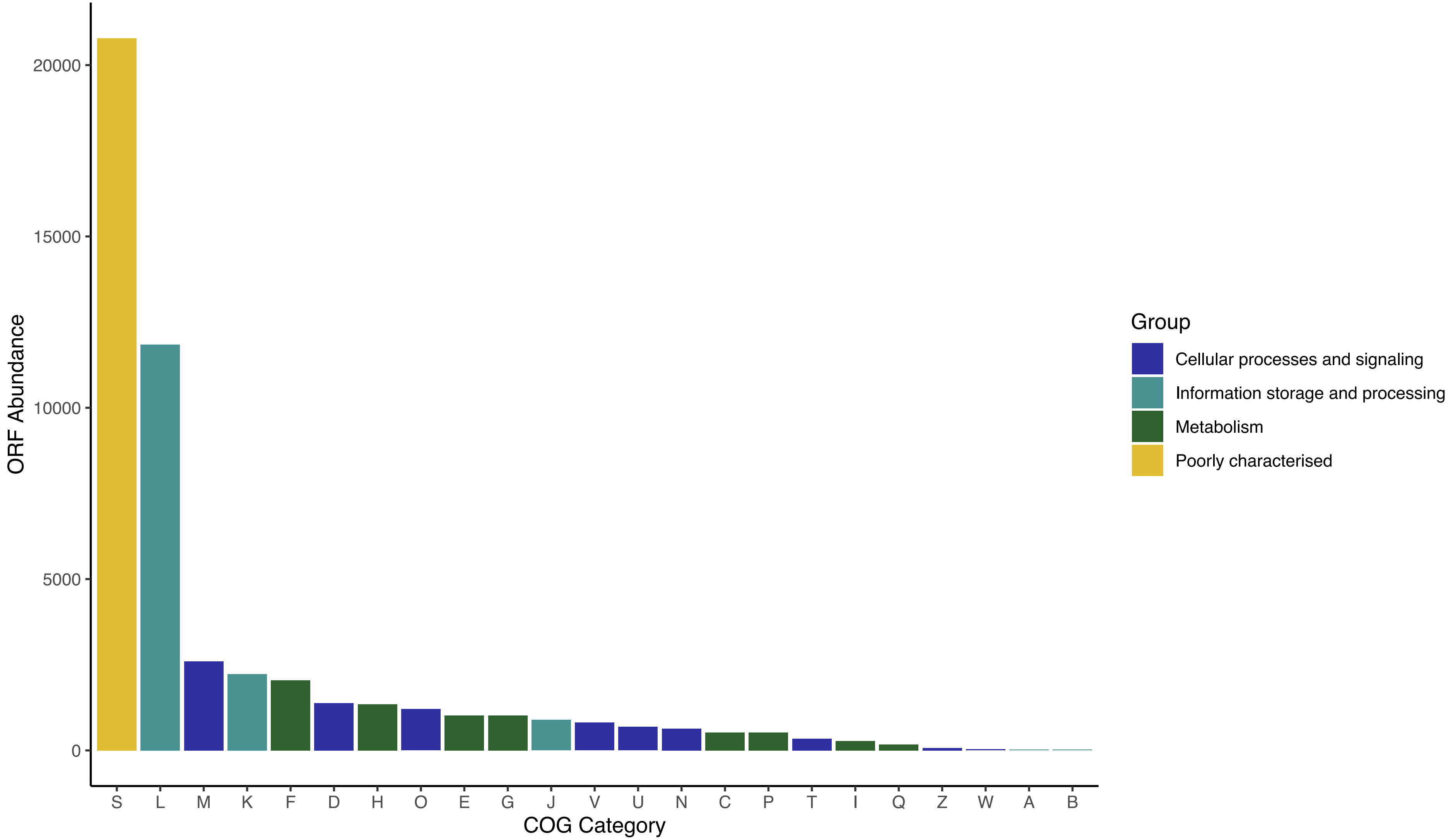

Supplementary figure 5

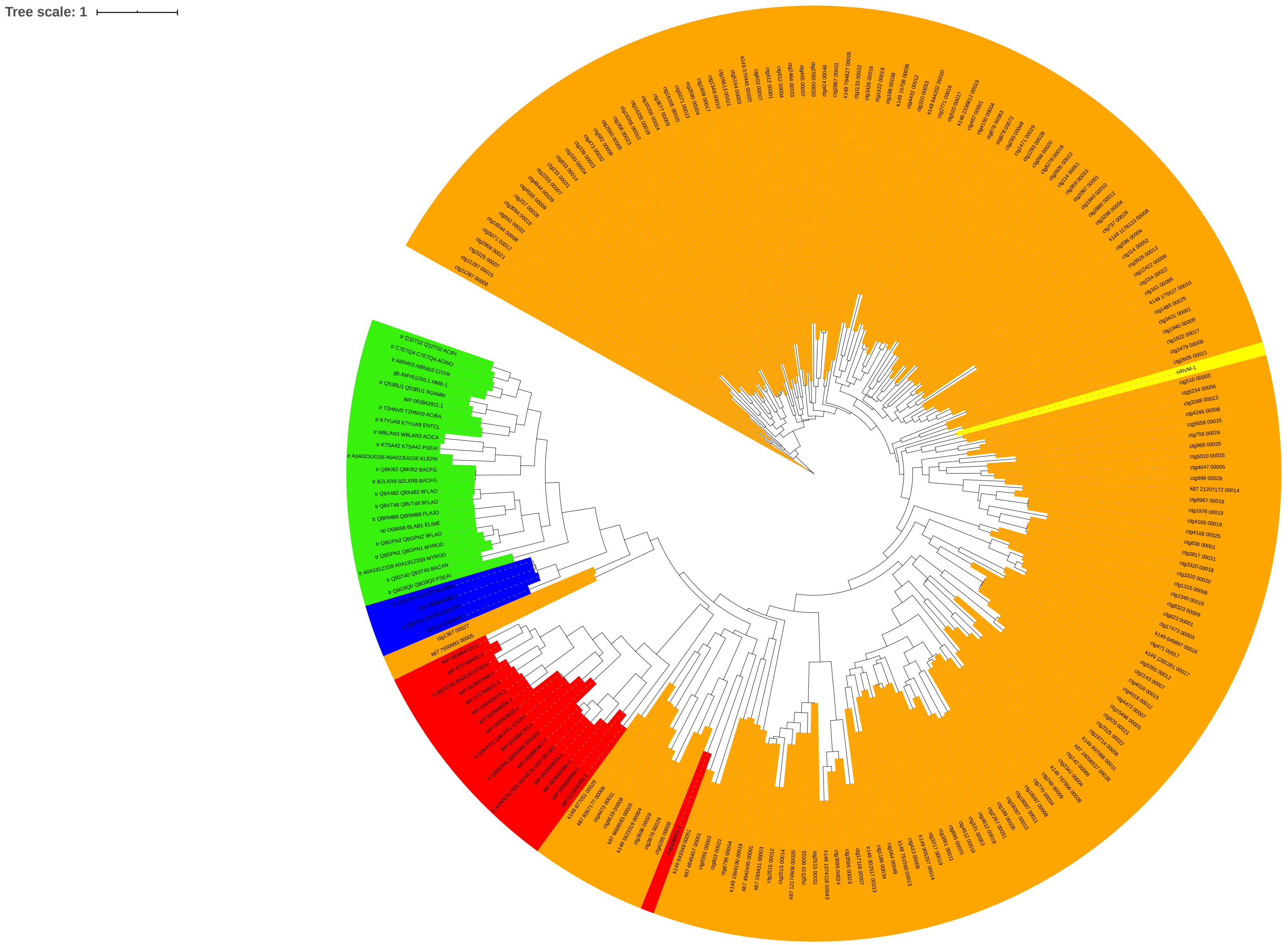

Supplementary figure 6

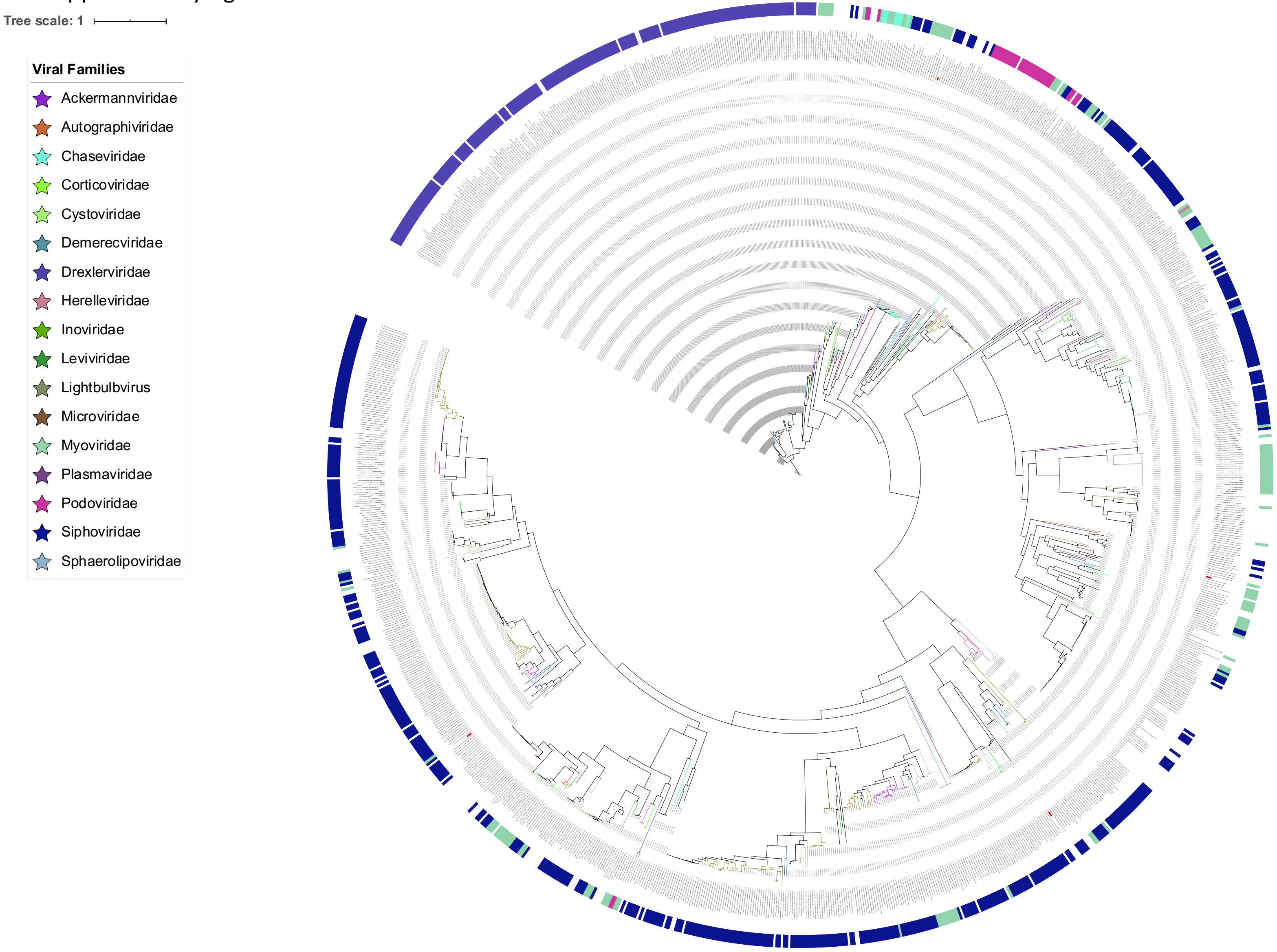
